# Human Striatal Association Megaclusters

**DOI:** 10.1101/2023.10.03.560666

**Authors:** Heather L. Kosakowski, Noam Saadon-Grosman, Jingnan Du, Mark E. Eldaief, Randy L. Buckner

## Abstract

The striatum receives projections from multiple regions of the cerebral cortex consistent with its role in diverse motor, affective, and cognitive functions. Supporting cognitive functions, the caudate receives projections from cortical association regions. Building on recent insights about the details of how multiple cortical networks are specialized for distinct aspects of higher-order cognition, we revisited caudate organization using within-individual precision neuroimaging (n=2, each participant scanned 31 times). Detailed analysis revealed that the caudate has side-by-side zones that are coupled to at least Give distinct distributed association networks, paralleling the specialization observed in the cerebral cortex. Examining correlation maps from closely juxtaposed seed regions in the caudate recapitulated the Give distinct cerebral networks including their multiple spatially distributed regions. These results extend the general notion of parallel specialized basal ganglia circuits, with the additional discovery that even within the caudate, there is Gine-grained separation of multiple distinct higher-order networks.

---

The basal ganglia support motor, affective, and cognitive functions through polysynaptic circuits that initiate in the cerebral cortex and return to cortex via the thalamus. Cortical projections to the striatum serve as the entry point for basal ganglia circuits (Glees 1944). Providing a substrate for influences on broad functional domains, widely distributed association regions, including multiple regions within prefrontal cortex (PFC), project to extended zones of the striatum (Goldman & Nauta 1977; Yeterian & Van Hoessen 1978; Selemon & Goldman-Rakic 1985). In a landmark synthesis, Alexander, DeLong, and Strick (1986) surmised that the basal ganglia support multiple distinct functions through specialized parallel closed-loop circuits. They proposed five circuits that each maintain segregation within the basal ganglia and project back to distinct frontal territories, including circuits that could support cognitive functions.

Anterograde tracing studies in non-human primates specifically reveal that dorsolateral PFC (DLPFC) projects to a region in the head of the caudate extending through the internal capsule and into the ventral margin of the putamen (e.g., Selemon & Goldman-Rakic 1985; see also Choi et al. 2017 for additional convergent cases). One critical detail is that, depending on the specific region of PFC, the projection targets vary across the medial to lateral extent of the caudate, suggesting that there may be finer distinctions (Selemon & Goldman-Rakic 1985; Ferry et al. 2000; Korponay et al. 2020). However, fine-grained topographic differences between projections to the caudate have been difficult to resolve, and there are indications that distributed regions of PFC may converge in the striatum.

Dual tracer injections from frontal and parietal regions yield substantially overlapping (interdigitated) striatal projection patterns in some cases, and in other cases minimal overlap (Selemon & Goldman-Rakic 1985). In a thorough quantitative analysis of 34 tracer injections in macaque PFC, Averbeck et al. (2014) noted variation in the patterning of projections to the striatum as well as an overlapping zone in the medial caudate that receives projections from multiple distinct regions of PFC (see also Choi et al. 2017). Exploring whether multiple, independent networks associated with higher-order cognition have distinct representation within the caudate is a major focus of the present study. The caudate is small and the topography of the cortical inputs is incompletely understood, yet the distinctions among cortical association networks that have recently emerged provide an impetus to revisit striatal organization using precision within-individual approaches.

Providing a foundation for our work, human neuroimaging studies using functional connectivity MRI (fcMRI) reliably find that PFC networks include the caudate (e.g., Di Martino et al. 2008; Barnes et al. 2010; Choi et al. 2012; Greene et al. 2014; Jarbo & Verstynen 2015; Marquand et al. 2017; Gordon et al. 2022; O’Rawe & Leung 2022). For example, Choi et al. (2012) analyzed data from 1,000 individuals and found that the caudate was robustly correlated with networks that involve distributed PFC regions. However, fine distinctions between association networks were not apparent. For example, seed regions placed in a PFC network linked to putative cognitive control involving DLPFC and a region along the frontal midline involved in distinct functions displayed substantially overlapping patterns of correlation in the caudate (see also Choi et al. 2017). Such an observation is consistent with the possibility of convergence but also may simply be due to blurring that emerges when anatomical differences between individuals are averaged (see Greene et al. 2020 for discussion).

Gordon et al. (2022) recently explored striatal organization within intensively sampled individual participants (see also Greene et al. 2020). They observed patterns generally consistent with the prior group-averaged analyses but also resolved spatial details that were not evident in prior work. Critically, regions associated with anterior, dorsal, and ventral zones of the caudate showed partially non-overlapping patterns in PFC. Contrasting an anterior region with a dorsal region in an exemplar dataset specifiically separated networks putatively involved in language functions from others. Greene et al. (2020) noted separation in the caudate between regions linked to a PFC network implicated in cognitive control and a distinct zone associated with the network commonly known as the Default Network. In quantitative analyses of overlap and separation among individuals, Greene et al. (2020) further concluded the caudate possesses multiple network-specific zones. These findings represent a level of anatomical specificity not possible when averaging group data, that we build upon here.

The present work utilizes a precision within-individual approach to explore in detail the striatal topography linked to distributed higher-order association networks. We utilize three specifiic advances to pursue this work. First, we analyzed data from two individual participants who were scanned extensively (over 31 separate MRI sessions) to boost the signal-to-noise ratio (SNR) and allow for direct replications. Second, we register the images across sessions using a processing pipeline optimized for within-individual registration and minimization of ancillary spatial blurring (Braga et al. 2019). Finally, we base our explorations on the observation of spatially precise side-by-side juxtapositions of association networks across the cerebral cortex, specifiically focusing on five networks that have been replicated across multiple data samples and functionally dissociated from one another (Du et al. 2023). These five juxtaposed networks form Supra-Areal Association Megaclusters (SAAMs) within multiple zones of the cerebral cortex. Here we leverage these recent insights from precision explorations of the cerebral cortex to discover novel representations of multiple association networks within the human caudate.

## Methods

### Overview

The present work explored striatal organization via functional coupling to distinct distributed cortical networks. Within each of the two individuals studied, the data were divided into three separate within-individual data partitions allowing distinctions within the striatum, particularly the putative cognitive network zones, to be identified (Data Set 1) and then replicated (Data Set 2). Data Set 3 further allowed seed regions to be placed within the striatum itself to verify using another analysis approach that distinct, spatially specific distributed cerebral networks are associated with subregions of the caudate.

### Participants

Two native English-speaking right-handed adult women participated for payment (22-23 yrs; data previously reported in Braga et al. 2019; Xue et al. 2021; Du et al. 2023). Neither had a history of neurologic or psychiatric illness. Participants provided informed consent using protocols approved by the Institutional Review Board of Harvard University.

### MRI Data Acquisition

Data were acquired at the Harvard Center for Brain Science using a 3T Siemens Prisma-fit MRI scanner using the vendor-supplied 64-channel phased-array head-neck coil (Siemens Healthcare, Erlangen, Germany). Head motion was mitigated using foam and inflated padding. Participants were instructed to remain still, awake, and look at a rear-projected display through a custom-built mirror attached to the head coil. During BOLD scanning, participants fixated a centrally presented plus sign (black on a gray background). The scanner room was illuminated. Eyes were video recorded using an Eyelink 1000 Plus with Long-Range Mount (SR Research, Ottawa, Ontario, Canada), and alertness was scored during each functional run.

Each person participated in 31 MRI sessions over 28-40 wks with no sessions on successive days. Each session involved multiple resting-state fixation runs acquired using blood oxygenation level-dependent (BOLD) contrast (Kwong et al. 1992; Ogawa et al. 1992). A custom multiband gradient-echo echo-planar pulse sequence developed by the Center for Magnetic Resonance Research (CMRR) at the University of Minnesota was used (Xu et al. 2012; Xu et al. 2013; Van Essen et al. 2013; see also Setsompop et al. 2012): voxel size = 2.4 mm, repetition time (TR) = 1s, echo time (TE) = 32.6 ms, flip-angle = 64°, matrix 88 x 88 x 65, anterior-to-posterior (AP) phase encoding, multislice 5x acceleration. Slice positioning was automated (van der Kouwe et al. 2005), and signal dropout was minimized by selecting a slice 25° from the anterior-posterior commissural plane toward the coronal plane (Weiskopf et al. 2006; Mennes et al. 2014). Each run lasted 7 min 2 sec (422 frames with 12 frames removed for T1 equilibration). A dual-gradient-echo B0 fieldmap was acquired to correct for spatial distortions: TE = 4.45 and 6.91 ms with slice prescription / resolution matched to the BOLD sequence. A rapid T1w structural scan was obtained using a multi-echo magnetization prepared rapid acquisition gradient echo (ME-MPRAGE) three-dimensional sequence (van der Kouwe et al. 2008): voxel size = 1.2 mm, TR = 2.20 s, TE = 1.57, 3.39, 5.21, 7.03 ms, TI = 1,100 ms, flip-angle = 7°, matrix 192 x 192 x 176, in-plane generalized auto-calibrating partial parallel acquisition (GRAPPA) acceleration = 4.

### Exclusion Criteria and Quality Control

BOLD runs were screened for quality. Exclusion criteria included: 1) maximum absolute motion > 2 mm and 2) slice-based SNR < 130. S1 had 62 and S2 had 61 usable runs of data.

### Data Processing and Registration that Minimizes Spatial Blurring

Data were processed using an in-house preprocessing pipeline (“iProc”) that preserved spatial details by minimizing blurring and multiple interpolations (described in detail in Braga et al. 2019). The pre-processed data were taken directly from Xue et al. (2021), additionally processed using the 15-network cerebral network estimates reported in Du et al. (2023).

Briefly, data were interpolated to a 1-mm isotropic native-space atlas (with all processing steps composed into a single interpolation) that was then projected using FreeSurfer v6.0.0 to the fsaverage6 cortical surface (40,962 vertices per hemisphere; Fischl et al. 1999; Fischl 2012). Five transformation matrices were calculated: (1) a motion correction matrix for each volume to the run’s middle volume [linear registration, 6 degrees of freedom (DOF); MCFLIRT, FSL], (2) a matrix for field-map-unwarping the run’s middle volume, correcting for field inhomogeneities caused by susceptibility gradients (FUGUE, FSL), (3) a matrix for registering the field-map-unwarped middle BOLD volume to the within-individual mean BOLD template (12 DOF; FLIRT, FSL), (4) a matrix for registering the mean BOLD template to the participant’s T1w native-space image (6 DOF; using boundary-based registration, FreeSurfer), and (5) a non-linear transformation to MNI space (nonlinear registration; FNIRT, FSL). The individual-specific mean BOLD template was created by averaging all field-map-unwarped middle volumes after being registered to an up-sampled 1.2 mm and unwarped mid-volume template (interim target, selected from a low motion run acquired close to a field map). The T1w native-space template was then resampled to 1.0 mm isotropic resolution.

Confounding variables including 6 head motion parameters, whole-brain, ventricular signal, deep cerebral white matter signal, and their temporal derivatives were calculated from the BOLD data in T1w native space. The signals were regressed out from the BOLD data using 3dTproject, AFNI (Cox 1996, 2012). The residual BOLD data were then bandpass filtered at 0.01–0.10-Hz using 3dBandpass, AFNI (Cox 1996, 2012). For surface analyses, the native space data were resampled to the fsaverage6 standardized cortical surface mesh using trilinear interpolation (featuring 40,962 vertices per hemisphere; Fischl et al. 1999; Fischl 2012) and then surface-smoothed using a 2-mm full-width-at-half-maximum (FWHM) Gaussian kernel. For striatal analyses, MNI volume space data were smoothed using a 4-mm FWHM Gaussian kernel. The iProc pipeline thus allowed for robustly aligned BOLD data, with interpolation minimized. Relevant final output spaces included the MNI volume space and the fsaverage6 cortical surface.

### Temporal Signal-to-Noise Ratio (tSNR) Maps

Data using BOLD-contrast (T2* images) and echo-planar imaging result in variable distortion and signal dropout due to magnetic susceptibility artifacts (e.g., Ojemann et al. 1997). Voxelwise temporal signal-to-noise ratio (tSNR) maps were computed for each participant. tSNR was calculated for each voxel in the volume by dividing the mean time course within an fMRI run by its standard deviation and then averaged across runs.

### Striatum IdentiIication and Visualization

A key step for the present inquiry was to ensure high-quality parcellations of the striatum within individual participants. To isolate striatal voxels, we identified the caudate, putamen, and nucleus accumbens in each participant using the FreeSurfer automated parcellation (Fischl 2012). The FreeSurfer-based masks were combined, binarized, and warped to MNI space using the same individual-specific matrix used in “iProc” preprocessing. Striatum masks were created for different purposes that included: (1) visualization of the striatal boundaries, (2) parcellation of the striatum allowing for voxel assignments that edge just past the T1w boundaries^1^, and (3) visualization of the cortex adjacent to the striatum. To accommodate the different uses of striatal masks, the boundary of the striatum was dilated at different levels using *fslmaths* and binarized. 1x dilation was used for striatal boundary visualization, 3x was used for striatal parcellation, and 5x dilation was used to visualize the adjacent cerebral cortex. The 5x dilation mask was specifically used in control checks to determine the effect of signal blurring of the cerebral cortex into the striatum. Striatum parcellations overlaid on individual T1w and mean T2* images are displayed in Figure 1.

**Figure 1.**
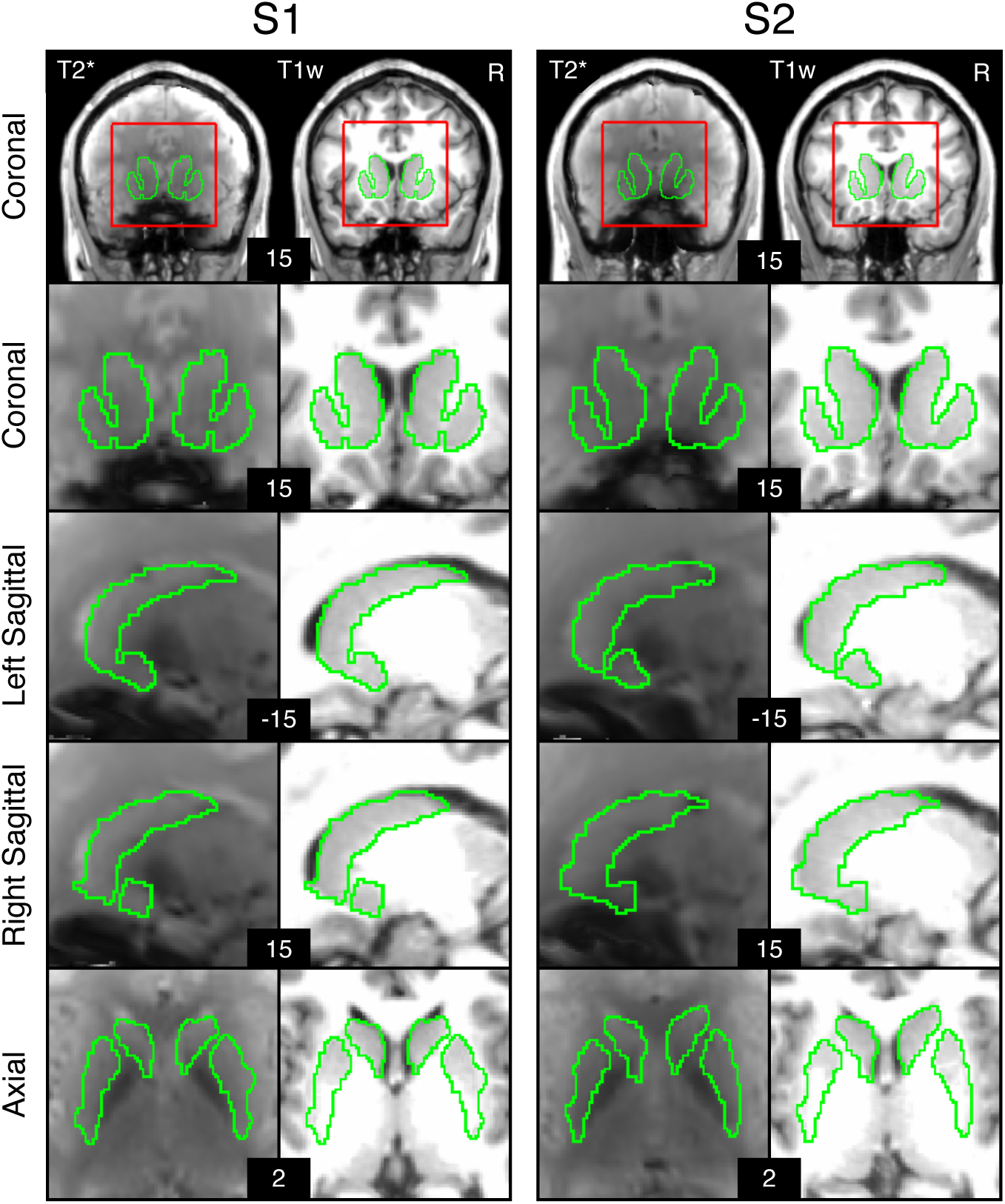
Automated identiGication of the striatum. Striatal boundaries in MNI atlas space are shown for S1 and S2. The first row depicts whole brain coronal slices. The red line indicates the bounding box used for striatal visualizations. The green line depicts the estimated boundaries of the striatum identified within each individual’s T1w structural image and then transformed to MNI atlas space. The estimated striatal boundaries are overlaid on T1w and average T2* images in registered MNI atlas space to illustrate distortion and signal drop – especially in the ventral striatum near the region of the susceptibility artifact.

### Individualized Network Estimates of the Cerebral Cortex

For both participants, the 15-network cerebral cortex estimates were taken directly from Du et al. (2023). A Multi-Session Hierarchical Bayesian Model (MS-HBM) was implemented to estimate the cortical networks (Kong et al. 2019). Data were split into three subsets. For S1, Data Set 1 included runs 1-20, Data Set 2 included runs 21-40, and Data Set 3 included runs 41-62. For S2, Data Set 1 included runs 1-20, Data Set 2 included runs 21-40, and Data Set 3 included runs 41-61. The MS-HBM was independently implemented on the first two subsets (Data Sets 1 and 2). Data Set 3 was set aside to allow for seed-region based correlation analysis to confirm network specificity.

To estimate networks in Data Sets 1 and 2, the connectivity profile of each vertex on the cortical surface was first estimated as its functional connectivity to 1,175 regions of interest (ROIs) that uniformly distributed across the fsaverage6 surface meshes (Yeo et al. 2011). For each run of data, the Pearson’s correlation coefficients between the fMRI time series at each vertex (40,962 vertices / hemisphere) and the 1,175 ROIs were computed. The resulting 40,962 x 1,175 correlation matrix per hemisphere was then binarized by keeping the top 10% of the correlations to obtain the functional connectivity profiles.

Next, the MS-HBM was initialized with a group-level parcellation estimated from the HCP S900 data release using the clustering algorithm from our previous study (Yeo et al. 2011). The group-level parcellations were used to initialize the expectation-maximization (EM) algorithm for estimating parameters in the MS-HBM. The goal of applying the model in this study was to obtain the best estimate of networks within each individual participant’s dataset, not to train parameters and apply them to unseen data from new participants (Kong et al. 2019).

The 15 network estimates include: Somatomotor-A (SMOT-A), Somatomotor-B (SMOT-B), Premotor-Posterior Parietal Rostral (PM-PPr), Cingulo-Opercular (CG-OP), Salience / Parietal Memory Network (SAL / PMN), Dorsal Attention-A (dATN-A), Dorsal Attention-B (dATN-B), Frontoparietal Network-A (FPN-A), Frontoparietal Network-B (FPN-B), Default Network-A (DN-A), Default Network-B (DN-B), Language (LANG), Visual Central (VIS-C), Visual Peripheral (VIS-P), and Auditory (AUD).

Here we focus on five higher-order association networks that are juxtaposed throughout the cerebral cortex: FPN-A, FPN-B, LANG, DN-B, and DN-A.

### Within-Individual Striatum-to-Cerebrum Correlation Matrices

Data were divided into three separate partitions. Data Sets 1 and 2 were used as discovery and replication datasets for the striatum parcellation, and Data Set 3 was set aside and used for the model-free seed-region based fcMRI analysis.

For each participant, the pair-wise *Pearson* correlation coefficients between the fMRI time courses at each surface vertex were calculated for each fixation fMRI run, yielding an 81,924 x 81,924 matrix (40,962 vertices / hemisphere). Separately, the pair-wise Pearson correlation coefficients between the fMRI time courses at each volume voxel within a 3x dilated striatum mask and each cortical vertex were calculated for each fixation fMRI run yielding an 68,713 x 81,924 matrix. The matrix was then Fisher *r*-to-*z* transformed and averaged across all runs to yield a single best estimate of the within-individual correlation matrix. This *z*-scored matrix was used to explore network organization. The mean correlation maps were assigned to a cortical and subcortical template combining left and right hemispheres of the fsaverage6 surface and subcortex around the striatum of the MNI156 volume into the CIFTI format to interactively explore correlation maps using the Connectome Workbench’s wb_view software (Marcus et al. 2011; Glasser et al. 2013). The colorbar scales of correlation maps were thresholded to highlight individual-specific anatomy using the Jet look-up table for visualization.

### Individual-SpeciIic Striatum Parcellation

To functionally parcellate the striatum in each individual participant, we adapted a winner-takes-all strategy (Choi et al. 2012; Xue et al. 2021). The objective of the parcellation was to assign each striatal voxel to its most strongly correlated cerebral network. The correlation strength between each voxel in the striatum and each vertex on the cortical surface was computed. Then, for each voxel in the striatum, we identified the 400 surface vertices that had the strongest correlation to that voxel. We estimated the percent of MS-HBM network assignments of the 400 surface vertices and assigned the voxel to the network with the highest proportion of network assignments.

While our focus was on five association networks (FPN-A, FPN-B, LANG, DN-B, and DN-A), it is important to note that the striatal voxels were not constrained to be assigned solely to these networks. Rather, voxels were permitted to be assigned to their most correlated network among all 15 of the cortical networks. That is, every voxel in the striatum had an opportunity to be assigned to any of the 15 estimated cerebral networks.

### Adjustment for Signal Blurring from Adjacent Cortex

The striatum is close to the cortical surface, particularly along the frontal midline and where the cortex of the insula folds into the Sylvian fissure. This proximity can cause erroneous assignments within the striatum that are due to signal blur from the surrounding cortical surface (see Choi et al. 2012 and Greene et al. 2014 for similar considerations). To understand the potential impact of signal blur, we conducted a series of control analyses.

In these control analyses, we dilated the striatal mask from each individual such that it extended beyond the anatomical boundary of the striatum into the adjacent cortical regions and ventricles. For this visualization, we removed correlation values below a specified threshold. Specifically, for the top 400 voxel-to-vertex correlations, if a correlation was below a pre-determined threshold (e.g., *r*(*z*) < 0.10), the vertex was excluded from the count. For example, if 100 of the 400 vertices had a correlation value below the threshold, only the 300 vertices with correlations above the threshold would be included to determine the assignment. If all voxel-to-vertex correlation values were below the threshold, the voxel did not receive a network assignment. Using a high correlation value as our threshold and a mask that extended into the adjacent cortical ribbon enabled us to visualize cortical signal bleed into the striatum. We tested thresholds *r*=0.01, and *r*=0.05-0.40 in increments of 0.05 across unsmoothed and smoothed data to visualize whether any of our interpreted observations were potentially impacted by spatial blur.

### Model-Free Seed-Region Based Exploration of Striatal Network Assignments

A winner-takes-all approach makes a strong assumption that striatal voxels are associated with a single cortical network and thus could bias estimates to detect segregation rather than convergence. For example, a striatal voxel correlated at a roughly similar level to three separate cortical networks will obligatorily be assigned to a single network hiding the critical detail that the voxel is correlated with a broad extent of the cerebral cortex (see also Greene et al. 2020). To provide another means to assess the specificity of networks linked to the striatum, the set aside Data Set 3 from each individual was used prospectively to test for network correlation patterns. This analysis provided for an independent confirmation of the network assignments using a distinct method in separate data.

Specifically, the parcellations from Data Sets 1 and 2 were used to guide seed-region placement. In the independent, left-out Data Set 3 from the same participant, seed regions were placed in zones of the caudate that belonged to FPN-A, FPN-B, LANG, DN-B, and DN-A. We then tested if the seed regions would recapitulate the distributed correlation pattern within the cerebral network boundaries of each network in the cerebral cortex.

As a complimentary analysis, we also explored the raw correlation patterns in the striatum from multiple seed regions placed in the cerebral cortex. Specifically, seed regions were placed in the MS-HBM-defined cerebral networks using the composite set of all fMRI data. For each network, we placed one seed region in four different spatially distributed nodes of the cerebral networks and observed correlation patterns in the striatum that corresponded to voxel assignments from the winner-takes-all parcellation strategy. The expectation is that raw correlation patterns within the striatum will be spatially diffuse given limitations of resolution but should also reveal spatial positioning differences consistent with the parcellation. Moreover, distinct cerebral seed regions from the same network should reveal convergent striatal patterns, and these convergent patterns should be distinct from the sets of seed regions positioned within distinct cerebral networks. That is, the raw correlation patterns of cerebral seed regions should reveal patterning in the striatum consistent with the automated network parcellation.

### Software and Statistical Analysis

Functional connectivity was calculated in MATLAB (version 2019a; MathWorks, Natick, MA) using Pearson’s product moment correlations. FreeSurfer v6.0.0, FSL, and AFNI were used during data processing. The estimates of networks on the cortical surface were visualized in Connectome Workbench v1.3.2. Model-free seed-region confirmations were performed in Connectome Workbench v1.3.2. Network parcellation was performed using code from (Kong et al. 2019) on Github (https://github.com/ThomasYeoLab/CBIG/tree/master/stable_projects/brain_parcellation/Kong2019_MS HBM).

## Results

### Summary

The goal of the present study was to characterize the organization of the striatum in humans. To do so, we analyzed densely sampled fMRI data collected from individual participants (Xue et al. 2021) and parcellated the striatum using a winner-takes-all approach (Buckner et al. 2011; Choi et al. 2012).

Specifically, each striatal voxel was assigned to the individual-specific cerebral network with which it was most correlated. Importantly, we did not constrain voxel assignments; any of the 15 estimated networks in the cerebral cortex were eligible. Remarkably, we observed network assignments to five distinct association networks in subregions of the caudate, recapitulating segregation and network adjacencies observed in the cerebral cortex. The striatal patterns replicated in a second, independent dataset from the same participants, and model-free seed-region based analyses in a third dataset from each participant confirmed that the caudate subregions were preferentially linked to distinct, distributed association networks.

### Automated Structural Parcellation of the Striatum

For each participant, the caudate, putamen, and nucleus accumbens were isolated. Figure 1 illustrates the striatal boundaries in relation to the average BOLD (T2*) and T1w structural data for each individual. Due to the dense within-individual fMRI data sampling, there was sufficient signal in most regions of the striatum for thorough exploration. Signal dropout was present near to and within the nucleus accumbens.

### Striatal Parcellations Reveal Close Juxtaposition of Subregions Linked to Distinct Cerebral Association Networks

The first step in generating the striatal parcellation was to estimate the organization of networks within the cerebral cortex. Figures 2 and 3 illustrate the anatomical positions of the five association networks of interest here (Du et al. 2023 provide comprehensive visualizations of all 15 networks). Of interest, five networks were robustly identified (and replicated, Figure S1) with close juxtapositions repeatedly across the cerebral cortex. We refer to these clusters of juxtaposed cerebral networks as Supra-Areal Association Megaclusters or SAAMs (Du et al. 2023). The putative cognitive control networks Frontoparietal Network-A (FPN-A) and Frontoparietal Network-B (FPN-B) were next to each other and spatially juxtaposed with the trio of domain-specialized networks, Language (LANG), Default Network-B (DN-B), and Default Network-A (DN-A). While the exact boundaries for the SAAMs varied, the spatial juxtapositions were apparent in both participants and across multiple parietal, temporal, and prefrontal association zones.

**Figure 2.**
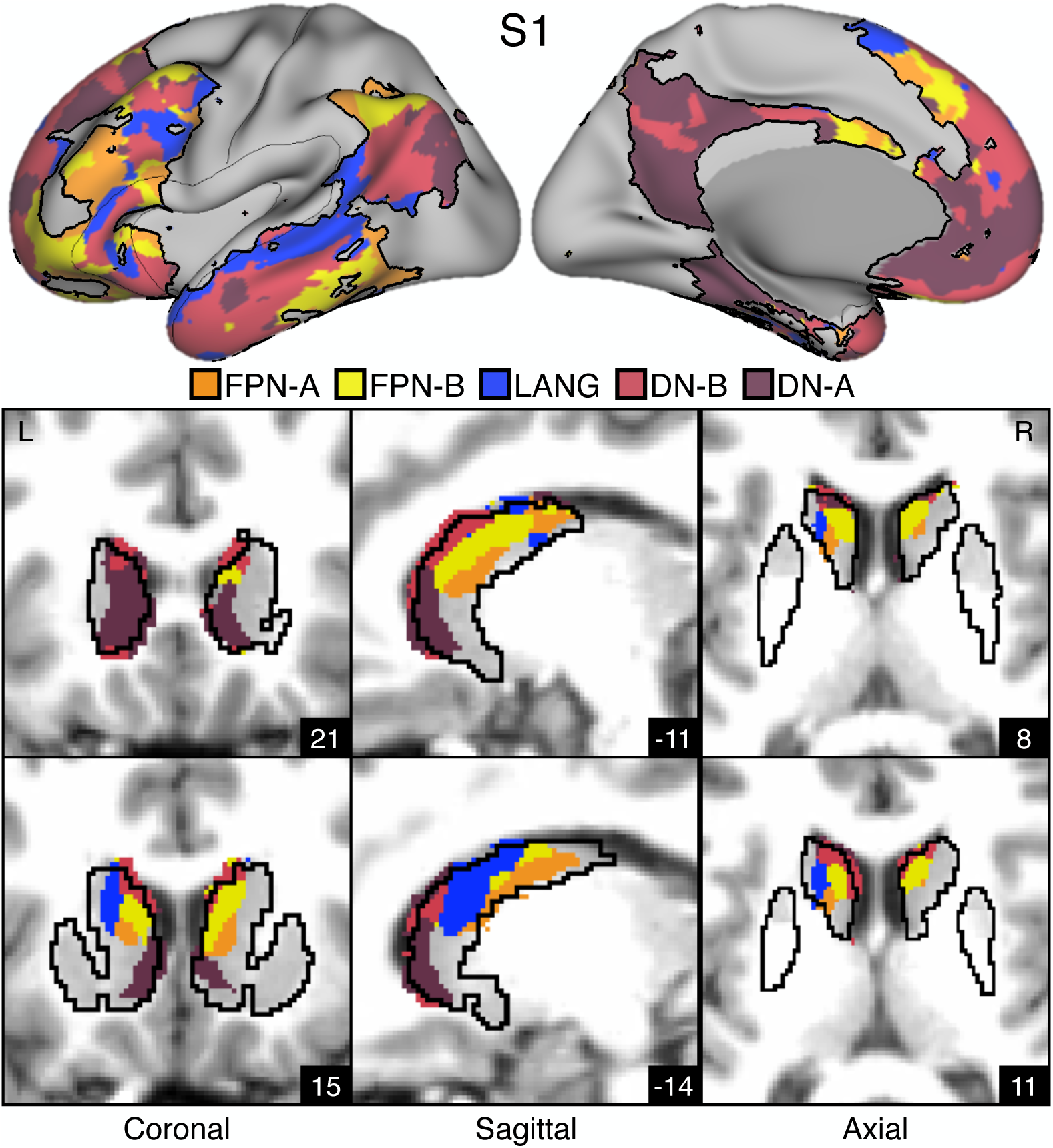
Striatal Association Megaclusters. Networks within the cerebral cortex were estimated including five higher-order association networks illustrated here (Du et al. 2023). The striatum was parcellated by assigning each voxel to its most correlated network in the cerebral cortex (see text). (**Top**) Inflated lateral and medial surfaces depict a cluster of five cortical networks that repeat across multiple cortical zones including posterior parietal cortex, temporal cortex, and prefrontal cortex (PFC). We refer to these repeating clusters as Supra-Areal Association Megclusters (SAAMs). Within each SAAM, FPN-A (orange), FPN-B (yellow), LANG (blue), DN-B (red), and DN-A (dark red) are adjacent to one another. The black outlines mark the combined borders around FPN-A, FPN-B, LANG, DN-B, and DN-A for several SAAMs. (**Bottom**) Multiple views of the striatum illustrate the representations of the five association networks in the caudate. Coordinates in the bottom right of each image indicate the slice in MNI152 space. Visualized data are from Data Set 1 for each participant. FPN-A, FPN-B, LANG, DN-B, and DN-A are observed with side-by-side relations within the caudate of both participants. These juxtapositions recapitulate the SAAMs within the cerebral cortex and are referred to as Striatal Association Megaclusters. See Supplementary Figure S2 for visualization of additional sections in both the discovery (Data Set 1) and replication (Data Set 2) datasets.

**Figure 3.**
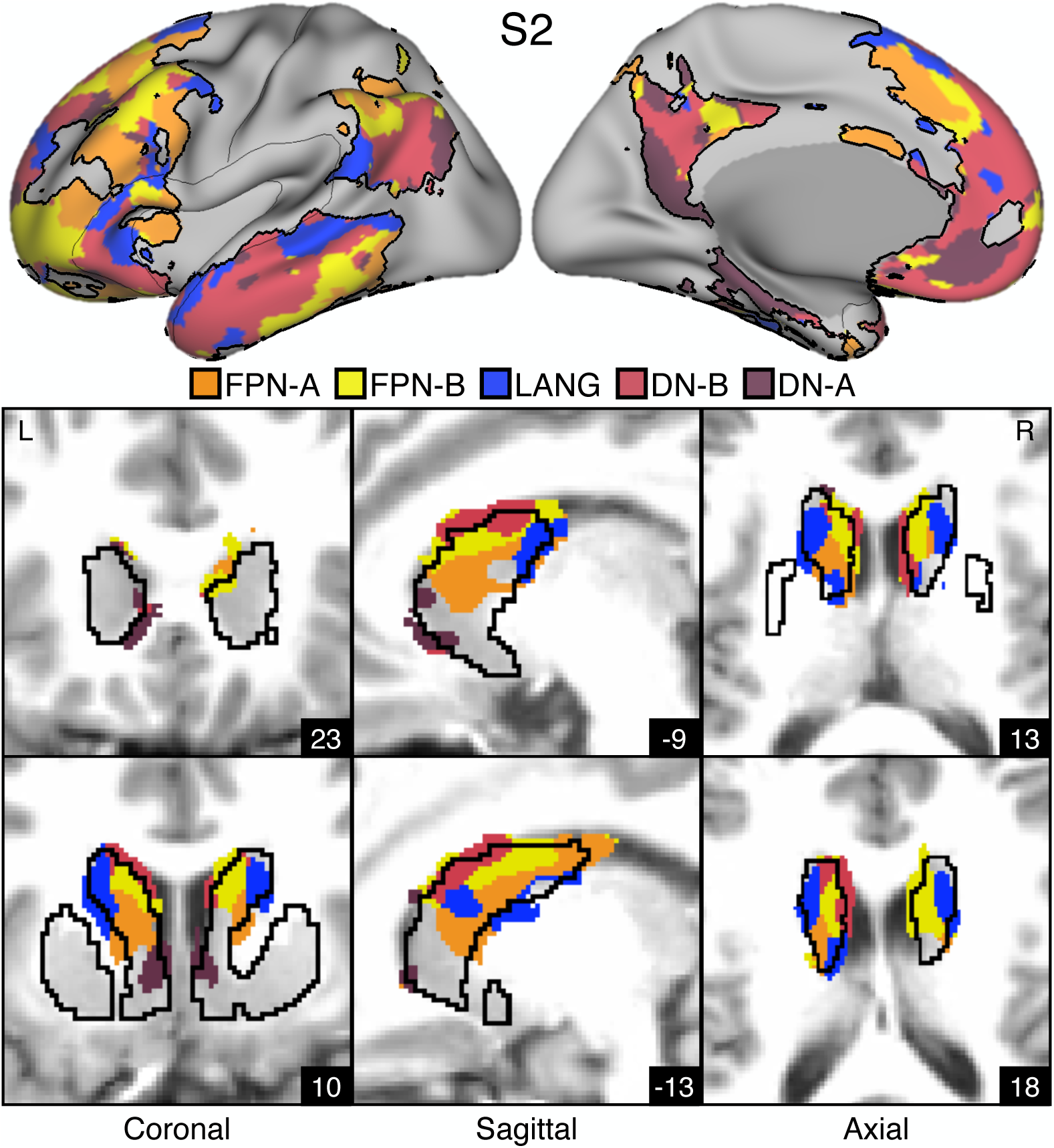
Striatal Association Megaclusters. Networks within the cerebral cortex were estimated including five higher-order association networks illustrated here (Du et al. 2023). The striatum was parcellated by assigning each voxel to its most correlated network in the cerebral cortex (see text). (**Top**) Inflated lateral and medial surfaces depict a cluster of five cortical networks that repeat across multiple cortical zones including posterior parietal cortex, temporal cortex, and prefrontal cortex (PFC). We refer to these repeating clusters as Supra-Areal Association Megclusters (SAAMs). Within each SAAM, FPN-A (orange), FPN-B (yellow), LANG (blue), DN-B (red), and DN-A (dark red) are adjacent to one another. The black outlines mark the combined borders around FPN-A, FPN-B, LANG, DN-B, and DN-A for several SAAMs. (**Bottom**) Multiple views of the striatum illustrate the representations of the five association networks in the caudate. Coordinates in the bottom right of each image indicate the slice in MNI152 space. Visualized data are from Data Set 1 for each participant. FPN-A, FPN-B, LANG, DN-B, and DN-A are observed with side-by-side relations within the caudate of both participants. These juxtapositions recapitulate the SAAMs within the cerebral cortex and are referred to as Striatal Association Megaclusters. See Supplementary Figure S2 for visualization of additional sections in both the discovery (Data Set 1) and replication (Data Set 2) datasets.

Striatal assignments to each of the five association networks revealed the presence of distinct subregions in the caudate in both participants that were linked to each of the five separate cerebral association networks (Figures 2, 3, S2). DN-A is found in the dorsomedial portion of the head of the caudate. DN-B surrounds DN-A but extends posteriorly from the head of the caudate through the body and into the tail. LANG is more lateral than DN-A and DN-B and crosses the anatomical boundaries of the caudate through the internal capsule and into the border of the putamen. Like DN-B, LANG begins in the head of the caudate and extends into the tail. Tightly interwoven FPN-A and FPN-B tend to be ventral to DN-B. Thus, the first major new result of our intensive within-individual analyses was that five distinct association networks remained segregated (or partially segregated) within the caudate. Moreover, the relative spatial positions within the caudate roughly recapitulated the organization in the cortex, with FPN-A and FPN-B subregions next to one another, surrounded by subregions associated with networks LANG, DN-B, and DN-A. For both participants, the spatial pattern of results was replicated in a second, independent dataset (Figure S2).

We refer to the striatal zones containing clustered juxtapositions of the five association networks as Striatal Association Megaclusters.

### Striatal Association Megaclusters Recapitulate Spatially Distinct Cerebral Networks

A key test of segregation of the caudate subregions was undertaken by examining whether closely juxtaposed seed regions placed within the caudate could recapitulate the spatially distinct cerebral networks. In independent data (Data Set 3), seed regions placed in the DN-A caudate zone from the discovery and replication parcellations (Data Sets 1 and 2) produced correlations that largely respected the boundaries of the MS-HBM-defined cortical FPN-A (Figures 4A & 5A). The same was true for seed regions placed in FPN-B (Figures 4B & 5B), LANG (Figures 4C & 5C), DN-B (Figures 4D & 5D), and DN-A (Figures 4E & 5E) striatal zones. The correlation patterns do show some blur into adjacent networks. In particular, the FPN-A seed region also generated correlations in adjacent FPN-B. Similarly, the DN-B seed region primarily recapitulated DN-B but portions of LANG were also implicated.

**Figure 4.**
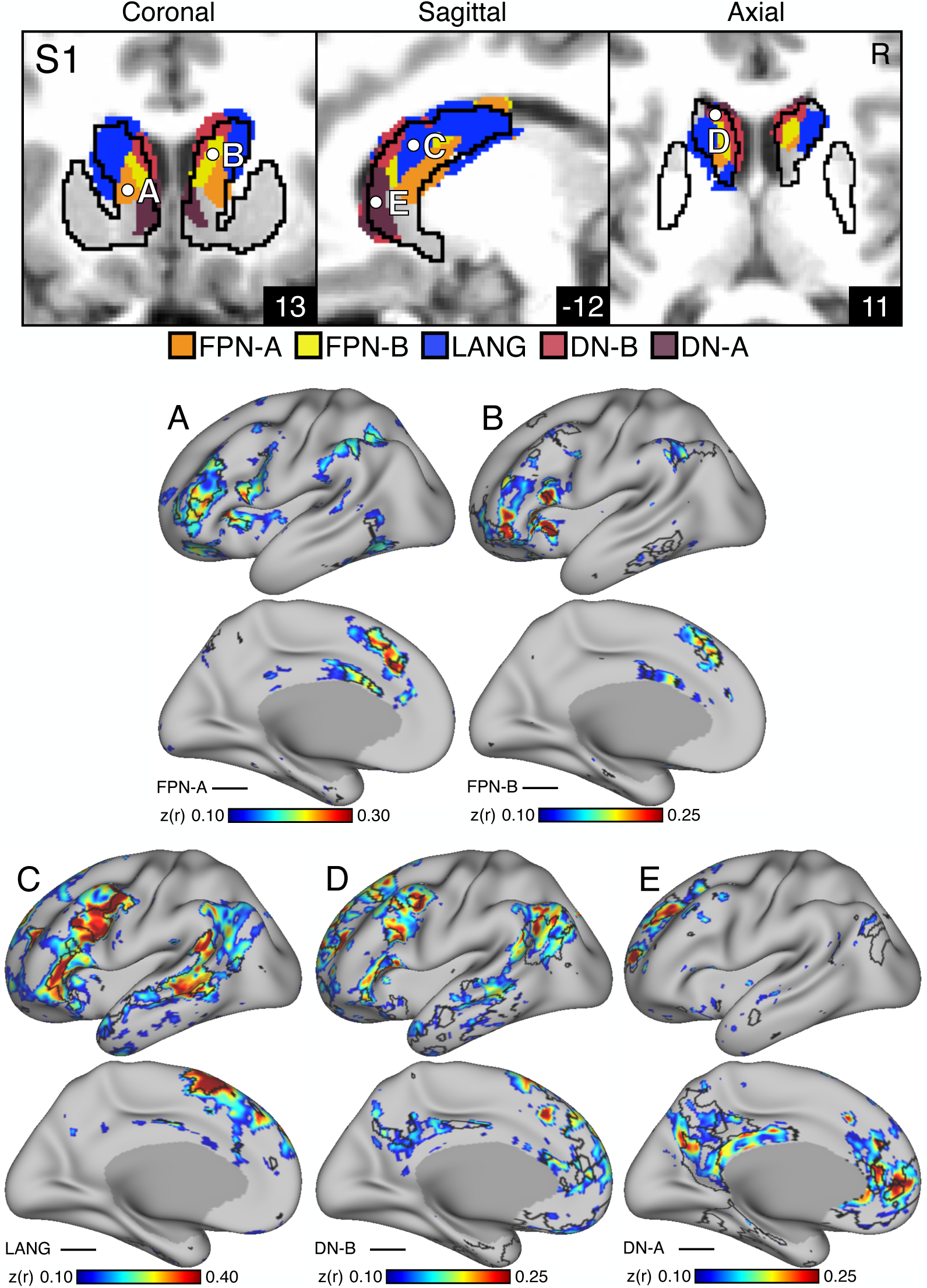
Subregions within the Striatal Association Megaclusters are correlated with parallel distributed association networks. To explore the specificity of subregions within the Striatal Association Megaclusters, a model-free seed-based fcMRI method was used to map cerebral networks correlated with side-by-side seed regions within the caudate. In each case, the striatum-to-cerebral cortex correlation maps are from independent data that were not used to derive the striatal parcellations (i.e., Data Sets 1 and 2 were used for striatal parcellations, Data Set 3 was used to estimate the striatum-to-cerebral cortex correlation maps). (**Top**) The striatal subregions associated with each network are shown in the caudate (Data Set 2 used for visualization) with white filled circles illustrating the locations of the five separate seed regions (labeled A to E). (**Bottom**) The seed region in the caudate assigned to FPN-A network (A) displays spatially selective correlation with the distributed cerebral FPN-A. Similarly, spatially selective cerebral networks are revealed for FPN-B (B), LANG (C), DN-B (D) and DN-A (E). The correlation maps are plotted as *z*(*r*) with the colorscale at the bottom. The Supplementary Materials display the patterns of striatal correlation from seed regions placed within the cerebral networks (Figures S5-S9 for S1 and Figures S10-S14 for S2).

**Figure 5.**
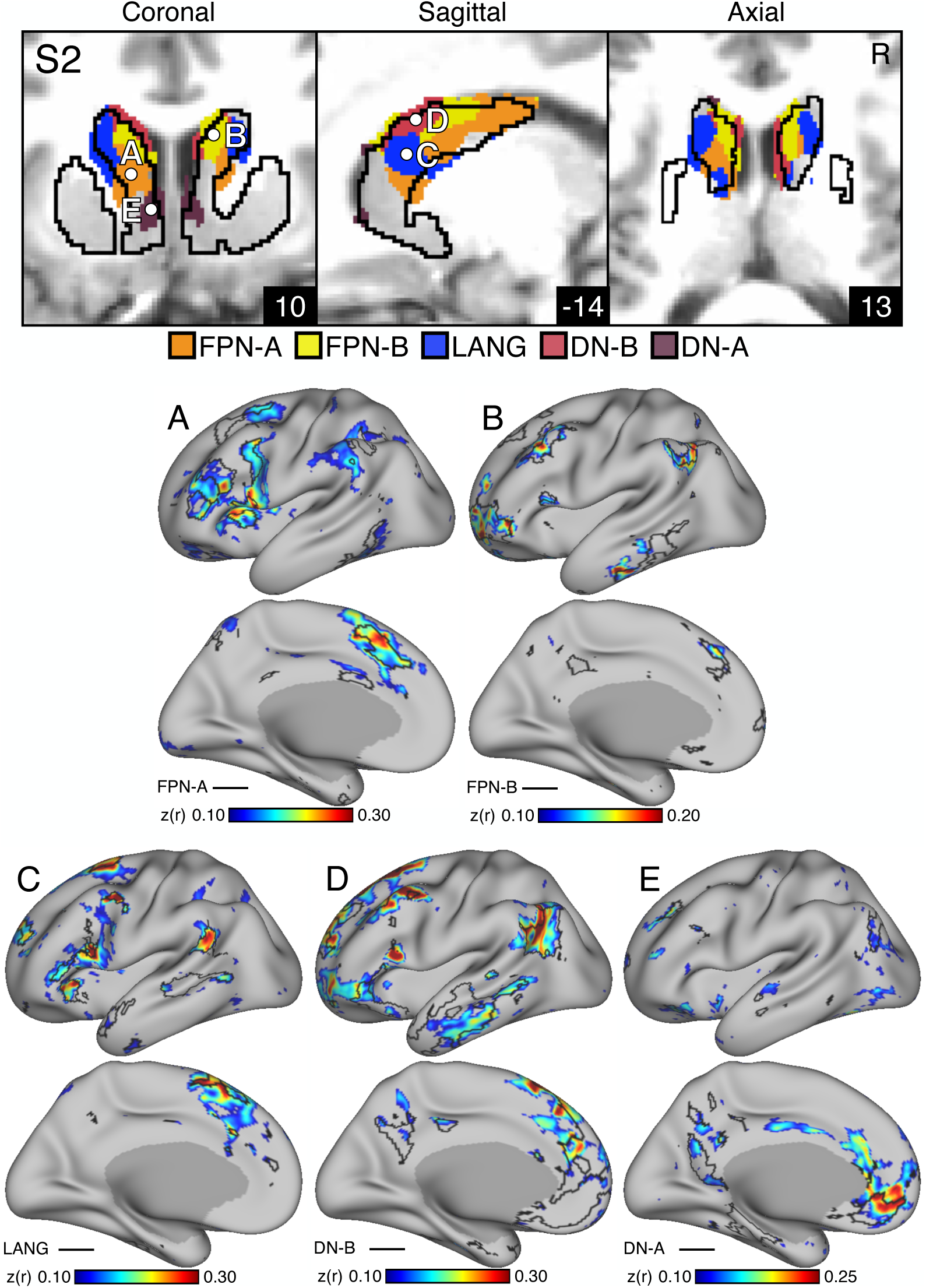
Subregions within the Striatal Association Megaclusters are correlated with parallel distributed association networks. To explore the specificity of subregions within the Striatal Association Megaclusters, a model-free seed-based fcMRI method was used to map cerebral networks correlated with side-by-side seed regions within the caudate. In each case, the striatum-to-cerebral cortex correlation maps are from independent data that were not used to derive the striatal parcellations (i.e., Data Sets 1 and 2 were used for striatal parcellations, Data Set 3 was used to estimate the striatum-to-cerebral cortex correlation maps). (**Top**) The striatal subregions associated with each network are shown in the caudate (Data Set 2 used for visualization) with white filled circles illustrating the locations of the five separate seed regions (labeled A to E). (**Bottom**) The seed region in the caudate assigned to FPN-A network (A) displays spatially selective correlation with the distributed cerebral FPN-A. Similarly, spatially selective cerebral networks are revealed for FPN-B (B), LANG (C), DN-B (D) and DN-A (E). The correlation maps are plotted as *z*(*r*) with the colorscale at the bottom. The Supplementary Materials display the patterns of striatal correlation from seed regions placed within the cerebral networks (Figures S5-S9 for S1 and Figures S10-S14 for S2).

### Cortical Signal Blurs into the Striatum

Although the replication of results within and across participants indicates the identified patterns are robust, there is a potential confound that could cause systematic errors in validity: cortical fMRI responses can be stronger than subcortical responses. Thus, it is possible that cortical signal blur could impact the parcellation within the striatum (see Choi et al. 2012 for discussion). This feature affects the multiple prior studies of striatal organization using human neuroimaging approaches including the present work. To test the impact of spatial blur from adjacent cortical regions, we used a winner-takes-all strategy on smoothed (4mm FWHM gaussian kernel) and unsmoothed data, where correlation values below a specific threshold were excluded. We systematically raised thresholds to observe if any voxel assignments in the striatum were continuations of cerebral correlation patterns. Figures S3 and S4 illustrate the results and indicate that signal blur from cortex may be impacting some striatal assignments.

Estimates in the lateral striatum, which borders insular cortex, and the ventral striatum, which borders orbitofrontal cortex, are ambiguous due to cortical signal bleed. Striatal Association Megaclusters in the caudate were largely (and clearly) separate from possible signal blur from adjacent cortex. One relevant striatal assignment of potential concern is the DN-A / DN-B cortical network portions that fall along the midline and could potentially extend into the striatum (see Figures S2 and S3, as well as the correlation maps in Figures S9 and S14). Thus, the location of the DN-A assignment in the caudate might be refined as higher resolution data are interrogated. Further, there are regions in more ventral portions of the striatum, which are not the focus of our results but were emphasized in our prior parcellations (Choi et al. 2012), that may be strongly affected by spatial blur given the proximity of the ventral regions of the striatum and the midline cortical frontal regions involved in DN-A and DN-B. Critically, the caudate zones associated with FPN-A, FPN-B, and LANG are particularly distinct by every measure (e.g., see the clear differences in even the raw correlations patterns in Figures S5 to S14) and discontinuous with any nearby cortical surface network assignments (Figures S3 and S4).

## Discussion

The basal ganglia operate on inputs to the striatum from diverse cortical regions with segregation between motor, affective, and cognitive loops (Alexander et al. 1986; see also Haber 2003). Here, using within-individual precision mapping of the striatum we discovered clustered representations of five separate higher-order association networks with side-by-side juxtapositions in the caudate. By placing separate seed regions within the five identified subregions of the caudate, we could recapitulate the topography of the distinct cerebral networks within each participant, supporting the inference that they participate in segregated or partially segregated networks. These results have conceptual implications for understanding striatal organization and function, as well as practical implications for the experimental study of the basal ganglia.

### Striatal Association Megaclusters

Separate representations of the five higher-order association networks were found within the caudate of both participants. We refer to these as Striatal Association Megaclusters because of their similarity to SAAMs present in the cerebral cortex, as reported by Du et al. (2023). Within the cerebral cortex, SAAMs are characterized by the juxtaposition of FPN-A, FPN-B, LANG, DN-B, and DN-A in a repeating fashion across parietal, temporal, and prefrontal zones. In the striatum, while there are idiosyncratic details that vary between participants, there are similar spatial relations among the five networks. DN-A is found in the dorsomedial portion of the head of the caudate with DN-B surrounding it and extending posteriorly through the body and into the tail. LANG is more lateral, begins in the head and extends into the anterior portion of the tail while FPN-A and FPN-B are interdigitated and ventral to DN-B. The relative spatial positions within the caudate recapitulate cortical organization, with FPN-A and FPN-B regions next to one another, surrounded by regions associated with networks LANG, DN-B, and DN-A. Importantly, this organization was not obligated by the analysis as voxels could be assigned to any one of the 15 cortical networks.

The presence of Striatal Association Megaclusters is consistent with the long-held idea that the basal ganglia possess multiple closed-loop circuits that operate on segregated channels of inputs (Alexander et al. 1986). Here, we revealed an unexpected level of specificity that segregates higher-order association networks within the caudate itself. This extension is surprising given prior neuroimaging studies (e.g., our own earlier work in Choi et al. 2012) and anatomical tract tracing studies that suggest convergence within the caudate (Averbeck et al. 2014). Our ability to detect and replicate spatially segregated subregions of the caudate is enabled by the precision within-individual approach adopted.

### Striatal Organization Parallels Cerebral and Cerebellar Organization

Following the advent of fcMRI to estimate cerebral networks (Biswal et al. 1995), within-individual precision methods have increased our understanding of the details of cortical organization (e.g., Laumann et al. 2015; Braga & Buckner 2017; Gordon et al. 2017; Braga et al. 2019, 2020; Smith et al. 2021; Somers et al. 2021; Noyce et al. 2022; Reznik et al. 2023; Gordon et al. 2023; Du et al. 2023; see also Fedorenko et al. 2010, 2011; Nieto-Castañ ón & Fedorenko 2012). Specifically, networks that initially appeared to involve large, less specialized regions were discovered in several instances to possess multiple tightly juxtaposed regions that are blurred together by group averaging methods (Laumann et al. 2015). For example, using precision approaches the extensively studied “Default Network” was found to be a combination of multiple, parallel networks with functionally dissociable side-by-side regions (Braga & Buckner 2017; Braga et al. 2020; DiNicola et al. 2020; Du et al. 2023; see Buckner & DiNicola, 2019 and DiNicola & Buckner, 2021 for discussion). In a recent discovery, precision mapping revealed the existence of inter-effector regions in the pre-central gyrus that are distinct from the canonical motor effector representations in M1 (Gordon et al. 2023). Precision mapping of the striatum, as reported here, similarly finds evidence for segregation in the caudate where prior work, including our own, found convergence (Choi et al. 2012; Choi et al. 2017).

In this light, it is interesting to emphasize that the Striatal Association Megaclusters parallel the complex yet elegant organization of cerebral and cerebellar association zones. In the cerebral cortex, multiple SAAMs with the same motif of network juxtapositions are present consistently across individuals (Du et al. 2023). The cerebellum possesses a parallel organization -- an association megacluster -- in at least one large zone. Specifically, in the cerebellum, there is a consistent clustering of the five networks around Crus I / II (see Xue et al. 2021 Figure 10). Further, Marek et al. (2018) used a distinct set of network estimates and revealed a similar set of side-by-side juxtapositions in Crus I / Crus II including distinctions between regions linked to FPN and DN. Thus, as precision mapping approaches have been applied, greater spatial specificity and segregation has been observed in both cerebral and cerebellar cortices. Our understanding of striatal organization has similarly evolved with the application of within-individual precision methods (see also Greene et al. 2020; Gordon et al. 2022).

The present findings are consistent with a framework that the basal ganglia and cerebellum are nodes in an integrated circuit with the cerebral cortex, and further that the integration involves separation of multiple functional domains (Bostan & Strick 2018). The repeating motif we find here in the striatum, and that was previously observed in the cerebellum, suggests that basal ganglia - cerebellar - cerebral cortical networks maintain segregated, or partially segregated, channels across multiple higher-order functional domains. This segregation forms megaclusters in the caudate that echo their cerebral counterparts and similar clusters in the Crus I / II apex of the cerebellum.

### Limitations and Open Questions

Here we relied on fcMRI analyses to infer network structure within the cerebral cortex and its relations to the striatum. There are a variety of caveats to such methods and their interpretation (Fox & Raichle 2007; Van Dijk et al. 2010; Murphy et al. 2013; Buckner 2013; Smith et al. 2013; Power et al. 2014; Xue et al. 2021). One specific limitation for estimating striatal organization is signal bleed from the cortex to the striatum. Voxels in the striatum that are spatially near insular and orbitofrontal cortices are the most likely to have corrupted signals impacted by stronger cortical signals. Therefore, it is important to carefully inspect striatal patterns in relation to nearby cortex. Figures S3, S4, S9, and S14 illustrate this challenge and reveal potential artifacts of cortical blur into the striatum in several zones that are not the focus of the present work (near to the ventral striatum in particular). Assignments to DN-A along the midline are potentially of concern and may require further refinement, a possibility that will likely require examination at a higher resolution and field strength (7T or beyond).

It is also important to note that, while we propose Striatal Association Megaclusters linked to five specific networks are an important organizational feature of the caudate, it does not negate the possibility that other cortical regions also project to the caudate. For example, extensive work with non-human primates indicates that the frontal eye field (FEF) projects to the caudate (Parthasarathy et al. 1992; Leichnetz 2001; Cui et al. 2003) and that the nearby supplementary eye field (SEF) has a different pattern of projections to a nearby region of the caudate (Shook et al. 1991; Parthasarathy et al. 1992). In preliminary analyses, we found an FEF-correlated region in the caudate that is spatially adjacent to the Striatal Association Megaclusters (Figure S15). Future work should acquire data to functionally localize FEF and SEF and measure how they are coupled with the striatum in humans along with SAAMs.

While we provide evidence for segregated or partially segregated regions, in terms of the macro-organization of the caudate, our findings do not imply that interactions are absent or that our methods are able to observe all local circuit features. For example, the dual injection cases of Selemon and Goldman-Rakic (1985) revealed that distinct cortical regions project to interdigitated terminal fields within the striatum. This feature was particularly prominent in Case 18 and suggests an anatomical feature of striatal organization that is beyond the present resolution of fMRI. Relatedly, some hypotheses about integration in the basal ganglia focus on how projection zones asymmetrically extend across anatomical domains. For example, Haber, Fudge, and McFarland (2000) examined projections to and from the midbrain using an extensive series of non-human primate tracer injections. They observed a hierarchy by which projections originating from the shell of the nucleus accumbens (NAc) extended into the loops of adjacent parallel circuits, forming a cascading influence on motor circuits (see also Haber & Knutson 2010). Our analyses of macro-organizational patterns in the striatum would not be able to detect such anatomical features.

The present results may also be relevant to interpreting certain puzzling macro-organizational features of anatomical projection patterns. Yeterian and Van Hoessen (1978) noted diverse projections to the caudate from distinct regions of association cortex. Comparing patterns across injections, they hypothesized that cerebral regions that are connected to one another also share projection territories in the striatum, consistent with the idea that they are nodes in distributed association networks (e.g., Goldman-Rakic 1988; Mesulam 1990). Selemon and Goldman-Rakic (1985) examined, in detail, paired injections across PFC and parietal association zones that showed some overlap but also, for certain injection pairs, clear separation in the caudate. They suggested that the connected cortical regions may not always converge on the same caudate zones, but rather be adjacent to one another (see also Selemon & Goldman-Rakic 1988). The present results are informative as to why such diversity in projection patterns might arise.

The cerebral association zones that possess distinct side-by-side regions are anatomically variable from one person to the next. The motif and relative positions are similar but the absolute positions of network regions on the cerebral surface are highly idiosyncratic (Du et al. 2023). If similar features are present in non-human primates, variable combinations of networks would be sampled from injection to injection. Consistent with this possibility, tracer injections across cases and association zones reveal label in the caudate that sometimes contain considerable macro-organizational overlap and in other instances side-by-side adjacencies (e.g., Yeterian & Van Hoessen 1978; Selemon & Goldman-Rakic 1985). Such patterns are expected if the injections are, by happenstance, hitting variable combinations of adjacent networks. It will be interesting, in the future, to chart whether comprehensive direct anatomical estimates in non-human primates converge with the indirect estimates reported here.

## Conclusions

We describe a new parcellation of the striatum that considers the idiosyncratic anatomical differences between people. In so doing, we observed Striatal Association Megaclusters of tightly juxtaposed zones of the caudate that are coupled with higher-order association networks FPN-A, FPN-B, LANG, DN-B and DN-A. These results extend the general notion of parallel specialized basal ganglia circuits (Alexander et al. 1986), with the additional discovery that there is fine-grained separation of multiple distinct networks, even within the caudate.

## Acknowledgments

We thank the Harvard Center for Brain Science neuroimaging core and FAS Division of Research Computing. We thank T. O’Keefe for assisting in optimization of data processing and R. Mair for MRI physics support. Many thanks to P. Angeli, L. DiNicola, W. Sun, S. Kaiser, J. Ladopoulou, and A. Xue for helpful conversations and in-house support. The multi-band EPI sequence was provided by the Center for Magnetic Resonance Research (CMRR) at the University of Minnesota.

## Figure Legends

**Figure S1.**
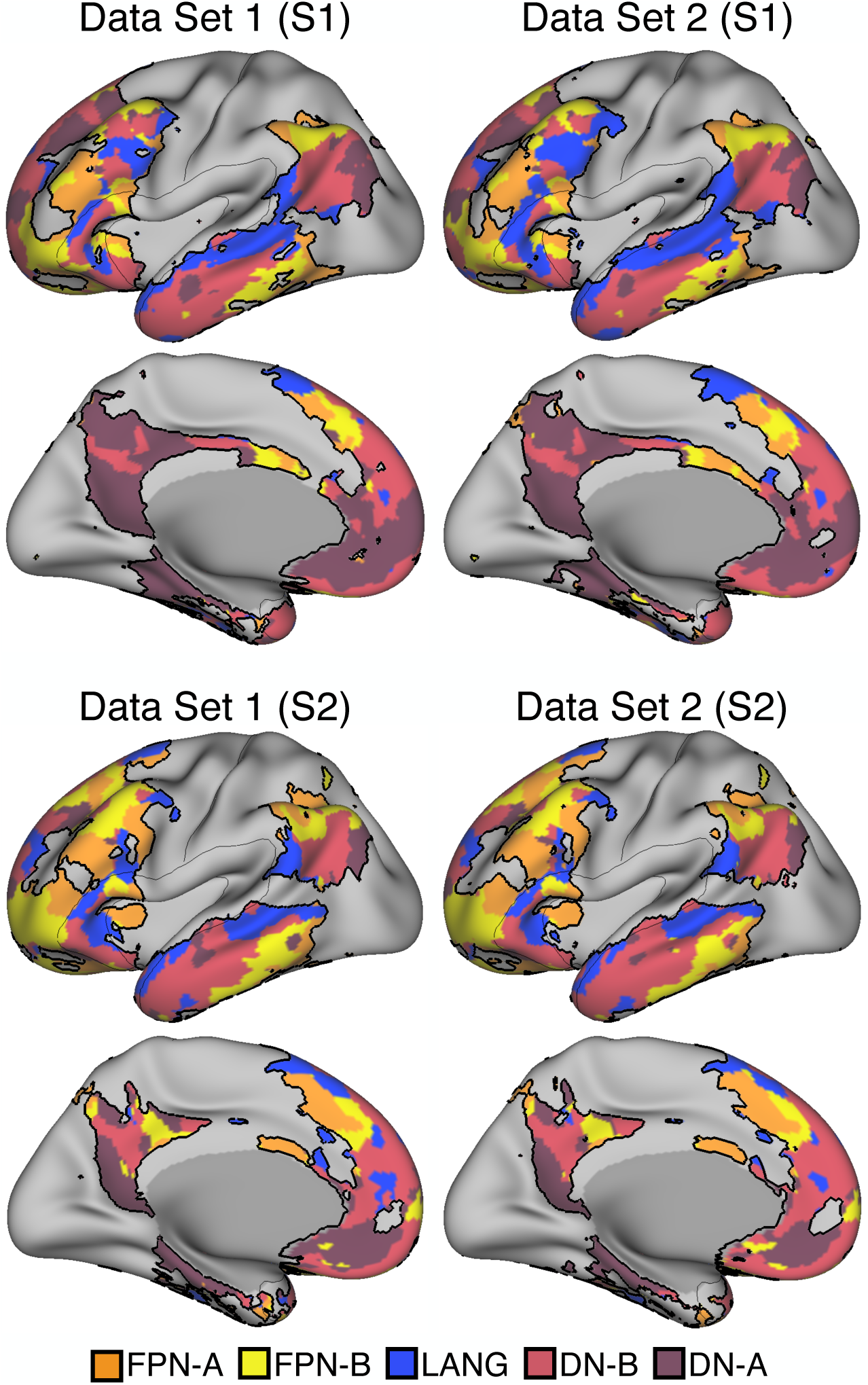
Cerebral Supra-Areal Association Megaclusters (SAAMs) are reliable across independent datasets within individuals. Independently analyzed subsets of data from S1 (left) and S2 (right) illustrate the reliability of FPN-A, FPN-B, LANG, DN-B, and DN-A network assignments. The juxtaposition and interdigitation of these networks are clearly evident and form Supra-Areal Association Megaclusters (SAAMs; outlined in black). The resting-state fixation data of S1 and S2 were split into three datasets (Data Sets 1, 2, and 3). The MS-HBM was applied to each dataset to independently estimate cortical networks (full network parcellation previously reported in Du et al. 2023). The individualized cortical network estimates replicate within participants. Network labels are shown at the bottom.

**Figure S2.**
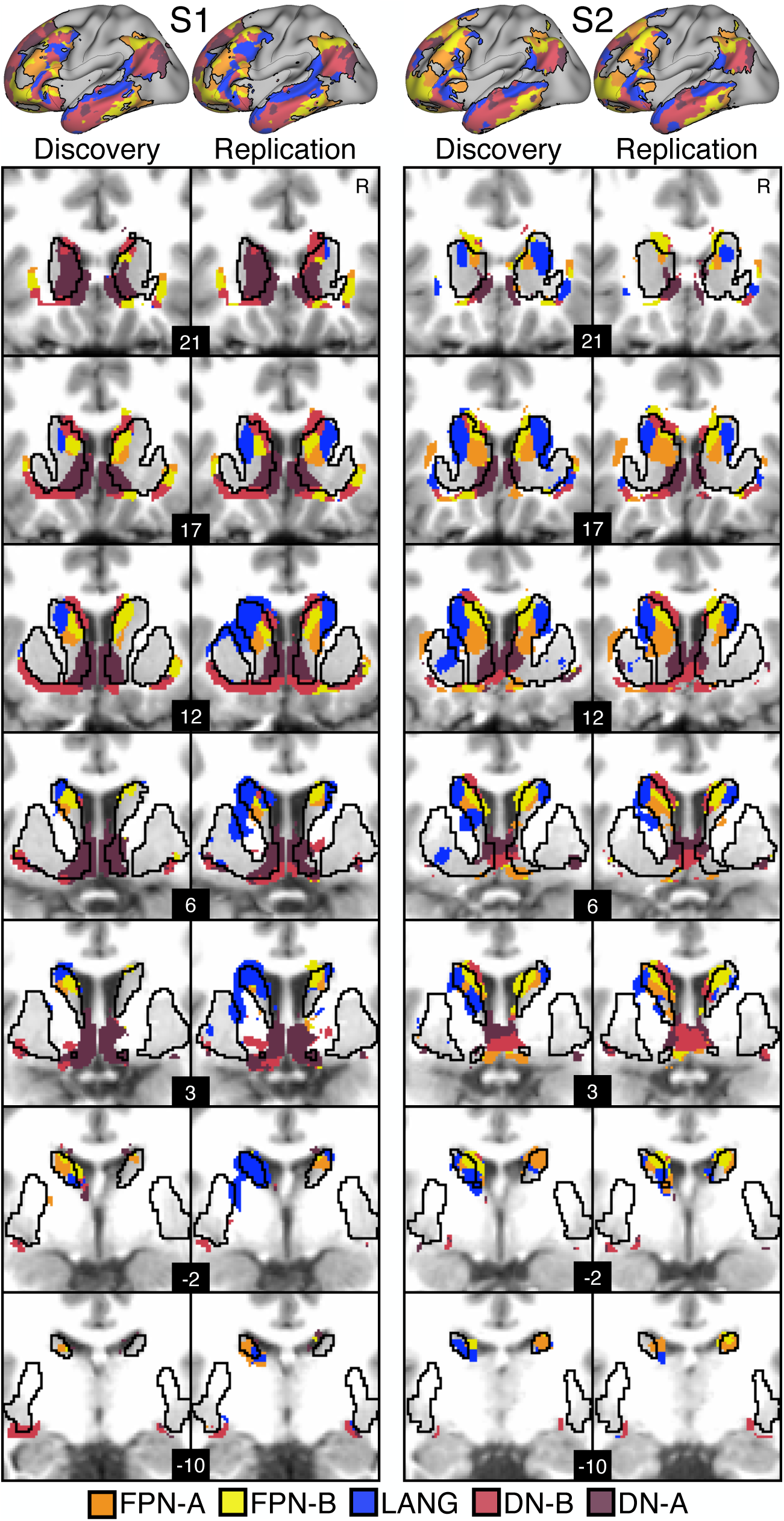
Striatal Association Megaclusters are reliable across independent datasets within individuals. An individual-specific winner-takes-all striatal parcellation strategy was independently applied to discovery (20 runs) and replication (20 runs) datasets. Each striatal voxel was assigned to one of fifteen cortical networks that were independently defined using a Multi-Session Hierarchical Bayesian Model (MS-HBM). Each voxel was assigned to the network that appeared the most often in the top 400 correlated vertices for that voxel. Shown here are all voxels that were assigned to association networks FPN-A (orange), FPN-B (yellow), LANG (blue), DN-B (red), and DN-A (dark red). Network assignments in the caudate are juxtaposed and clustered forming Striatal Association Megaclusters.

**Figure S3.**
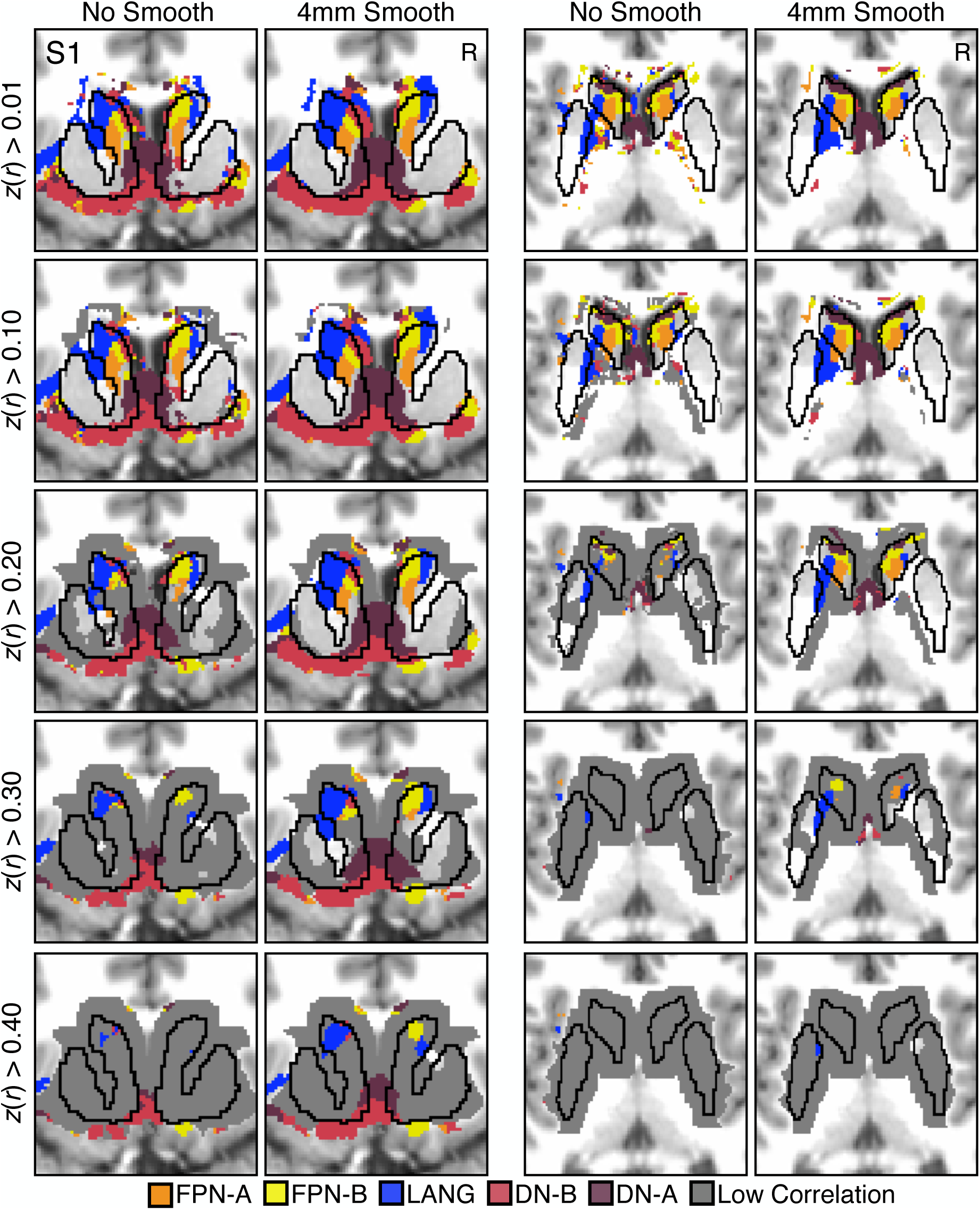
Signal bleed from cerebral cortex into lateral and ventral striatum. To optimize our striatal parcellation approach, we tested the effects of smoothing resting state fMRI data (no smooth and 4mm FWHM smoothing kernel) and excluding low correlation values from influencing network assignments. The striatum boundary was extended into the neighboring cortical area. For each voxel, we identified the top 400 vertex correlations. Any voxel-to-vertex correlation value (*z*(*r*)) that was less than a specified threshold (0.01, 0.10, 0.20, 0.30, and 0.40) was excluded from the analysis. If all *z*(*r*) values were below threshold, that voxel was not assigned to a network (gray). Using unsmoothed data resulted in the exclusion of striatal voxels (e.g., *z*(*r*)<0.20) but allowed for careful visualization of the relative contribution of cortical versus striatal signal. Including only high correlations (e.g., *z*(*r*)>0.40) primarily maintained voxel assignments in the cerebral cortex, demonstrating that cortical areas near the ventral striatum and lateral putamen bleed into, and influence striatal signals.

**Figure S4.**
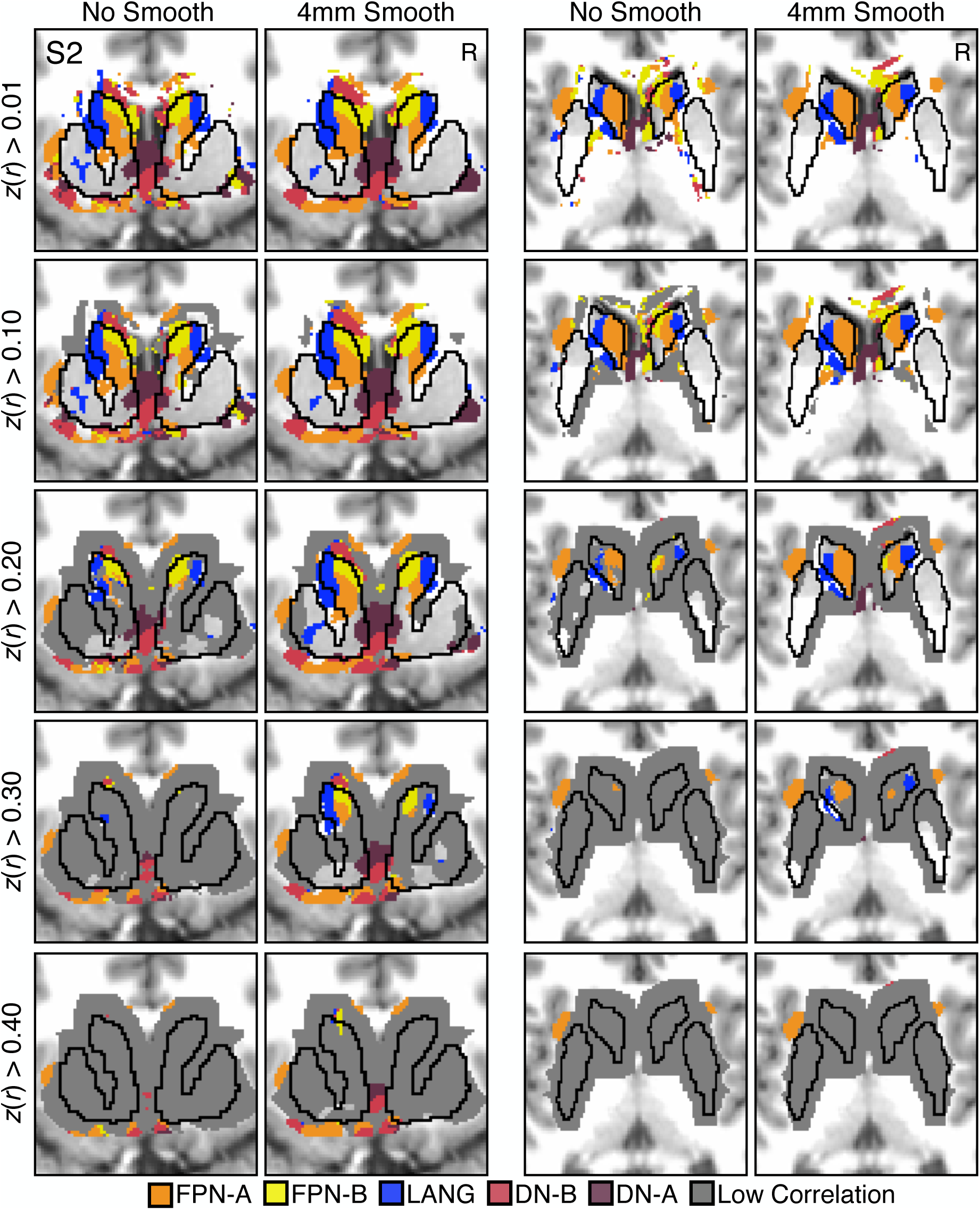
Signal bleed from cerebral cortex into lateral and ventral striatum. To optimize our striatal parcellation approach, we tested the effects of smoothing resting state fMRI data (no smooth and 4mm FWHM smoothing kernel) and excluding low correlation values from influencing network assignments. The striatum boundary was extended into the neighboring cortical area. For each voxel, we identified the top 400 vertex correlations. Any voxel-to-vertex correlation value (*z*(*r*)) that was less than a specified threshold (0.01, 0.10, 0.20, 0.30, and 0.40) was excluded from the analysis. If all *z*(*r*) values were below threshold, that voxel was not assigned to a network (gray). Using unsmoothed data resulted in the exclusion of striatal voxels (e.g., *z*(*r*)<0.20) but allowed for careful visualization of the relative contribution of cortical versus striatal signal. Including only high correlations (e.g., *z*(*r*)>0.40) primarily maintained voxel assignments in the cerebral cortex, demonstrating that cortical areas near the ventral striatum and lateral putamen bleed into, and influence striatal signals.

**Figure S5.**
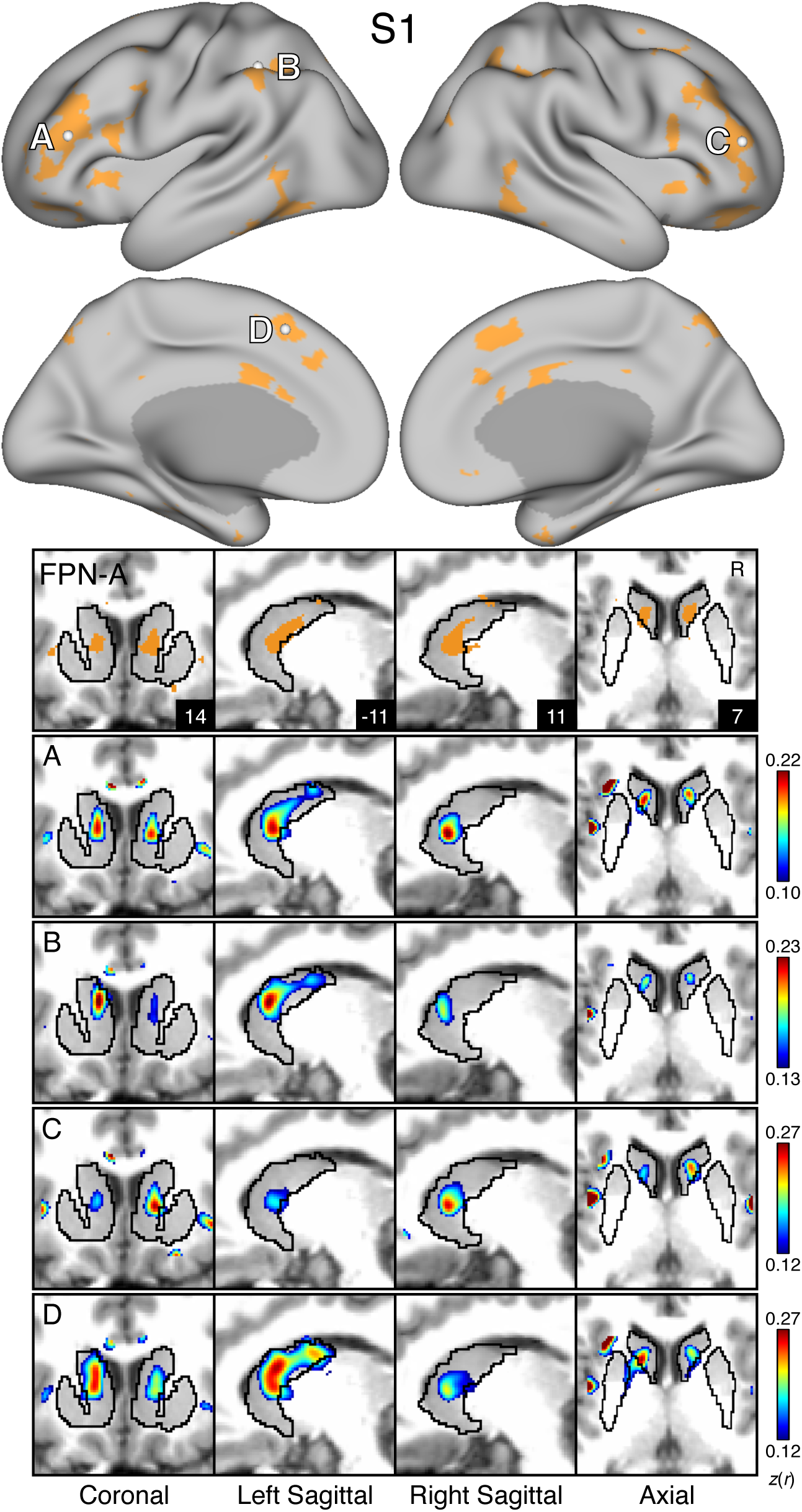
Model-free seed regions in cortical association networks recapitulate Striatal Association Megaclusters. For S1 and S2, the MS-HBM parcellation for a given network is shown on the cortex and in the striatum with the striatal boundaries outlined in black. Four seeds were placed in anterior and posterior nodes (A-D) of cortical association networks FPN-A (orange; Figures S5 and S10), FPN-B (yellow; Figures S6 and S11), LANG (blue; Figures S7 and S12), DN-B (red; Figures S8 and S13), and DN-A (dark red; Figures S9 and S14). Black outlines indicate the boundaries of corresponding individual-specific parcellation-defined networks estimated from the MS-HBM. The correlation maps are plotted as *z*(*r*) with the colorscale at the bottom. There is strong agreement between the seed-based correlation maps and the estimated network parcellation in the striatum.

**Figure S6.**
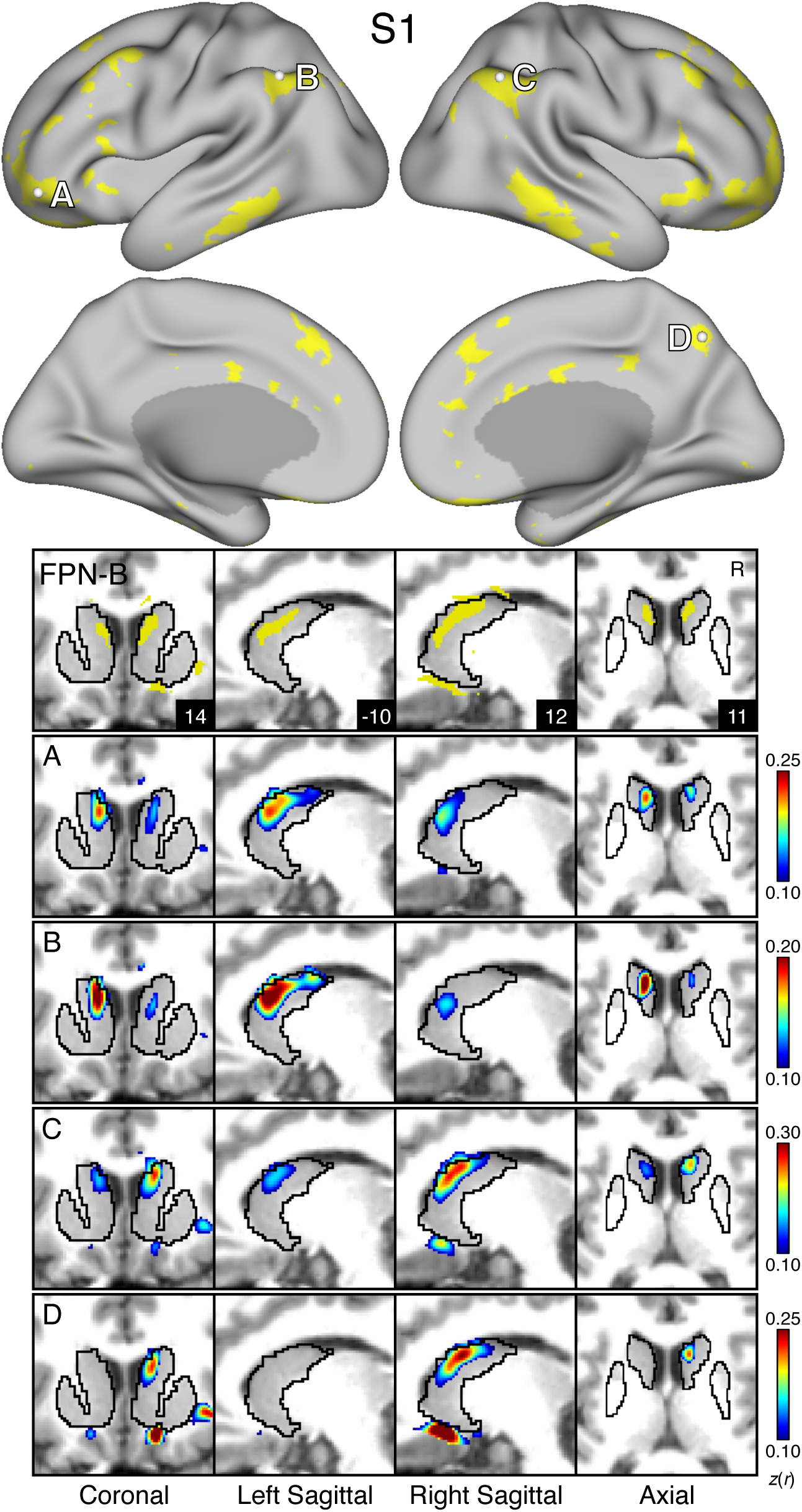
Model-free seed regions in cortical association networks recapitulate Striatal Association Megaclusters. For S1 and S2, the MS-HBM parcellation for a given network is shown on the cortex and in the striatum with the striatal boundaries outlined in black. Four seeds were placed in anterior and posterior nodes (A-D) of cortical association networks FPN-A (orange; Figures S5 and S10), FPN-B (yellow; Figures S6 and S11), LANG (blue; Figures S7 and S12), DN-B (red; Figures S8 and S13), and DN-A (dark red; Figures S9 and S14). Black outlines indicate the boundaries of corresponding individual-specific parcellation-defined networks estimated from the MS-HBM. The correlation maps are plotted as *z*(*r*) with the colorscale at the bottom. There is strong agreement between the seed-based correlation maps and the estimated network parcellation in the striatum.

**Figure S7.**
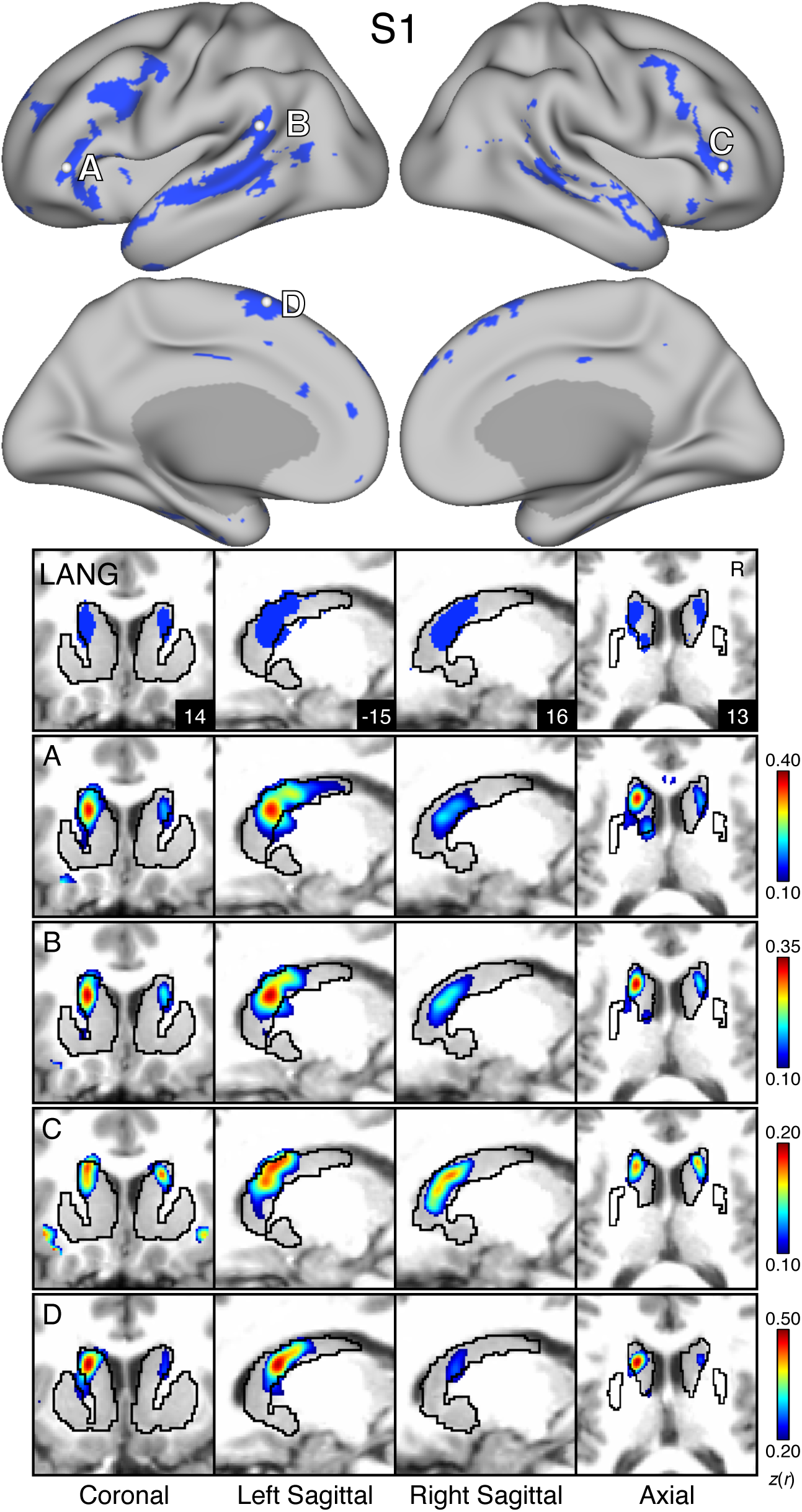
Model-free seed regions in cortical association networks recapitulate Striatal Association Megaclusters. For S1 and S2, the MS-HBM parcellation for a given network is shown on the cortex and in the striatum with the striatal boundaries outlined in black. Four seeds were placed in anterior and posterior nodes (A-D) of cortical association networks FPN-A (orange; Figures S5 and S10), FPN-B (yellow; Figures S6 and S11), LANG (blue; Figures S7 and S12), DN-B (red; Figures S8 and S13), and DN-A (dark red; Figures S9 and S14). Black outlines indicate the boundaries of corresponding individual-specific parcellation-defined networks estimated from the MS-HBM. The correlation maps are plotted as *z*(*r*) with the colorscale at the bottom. There is strong agreement between the seed-based correlation maps and the estimated network parcellation in the striatum.

**Figure S8.**
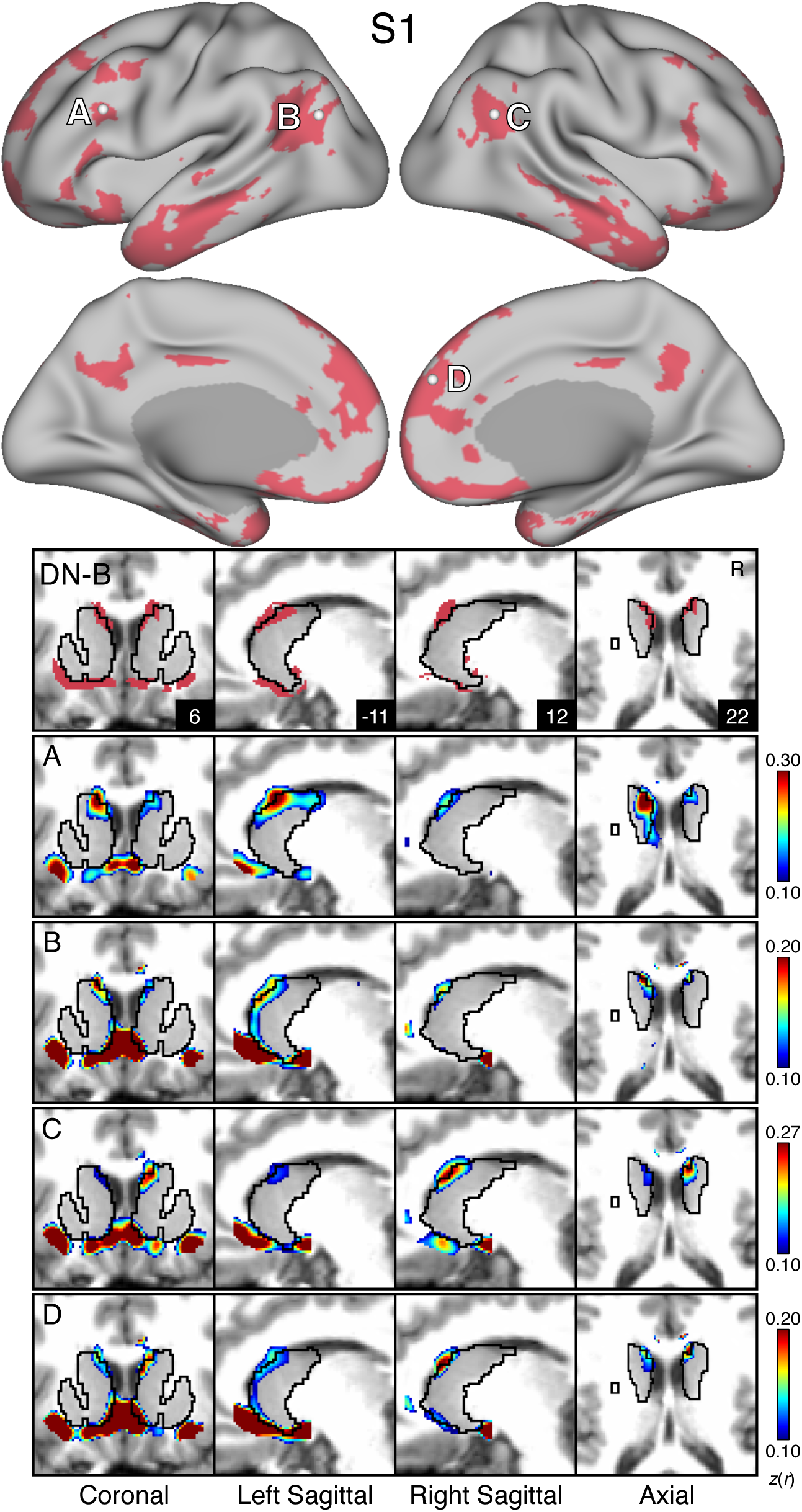
Model-free seed regions in cortical association networks recapitulate Striatal Association Megaclusters. For S1 and S2, the MS-HBM parcellation for a given network is shown on the cortex and in the striatum with the striatal boundaries outlined in black. Four seeds were placed in anterior and posterior nodes (A-D) of cortical association networks FPN-A (orange; Figures S5 and S10), FPN-B (yellow; Figures S6 and S11), LANG (blue; Figures S7 and S12), DN-B (red; Figures S8 and S13), and DN-A (dark red; Figures S9 and S14). Black outlines indicate the boundaries of corresponding individual-specific parcellation-defined networks estimated from the MS-HBM. The correlation maps are plotted as *z*(*r*) with the colorscale at the bottom. There is strong agreement between the seed-based correlation maps and the estimated network parcellation in the striatum.

**Figure S9.**
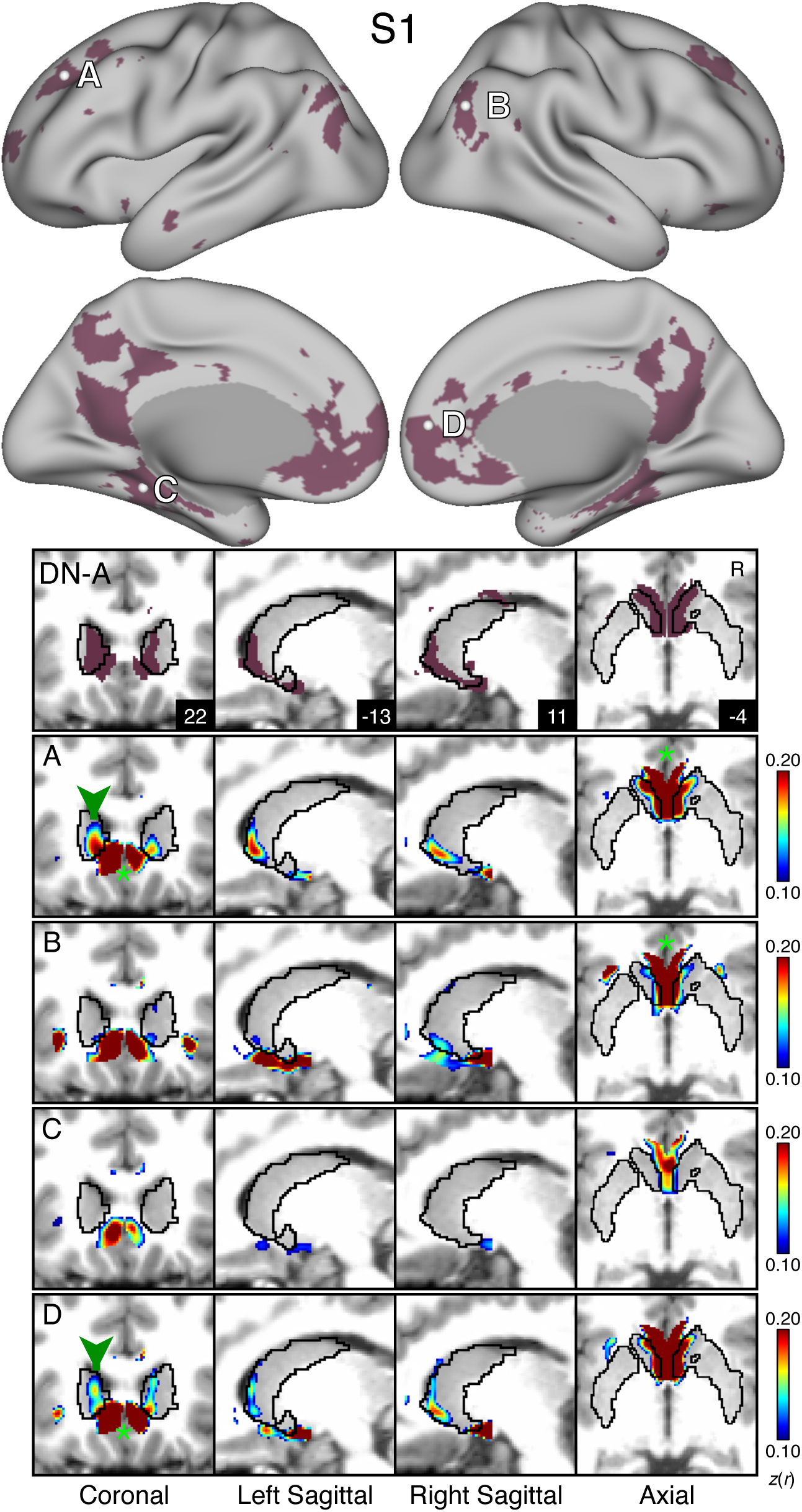
Model-free seed regions in cortical association networks recapitulate Striatal Association Megaclusters. For S1 and S2, the MS-HBM parcellation for a given network is shown on the cortex and in the striatum with the striatal boundaries outlined in black. Four seeds were placed in anterior and posterior nodes (A-D) of cortical association networks FPN-A (orange; Figures S5 and S10), FPN-B (yellow; Figures S6 and S11), LANG (blue; Figures S7 and S12), DN-B (red; Figures S8 and S13), and DN-A (dark red; Figures S9 and S14). Black outlines indicate the boundaries of corresponding individual-specific parcellation-defined networks estimated from the MS-HBM. The correlation maps are plotted as *z*(*r*) with the colorscale at the bottom. There is strong agreement between the seed-based correlation maps and the estimated network parcellation in the striatum.

**Figure S10.**
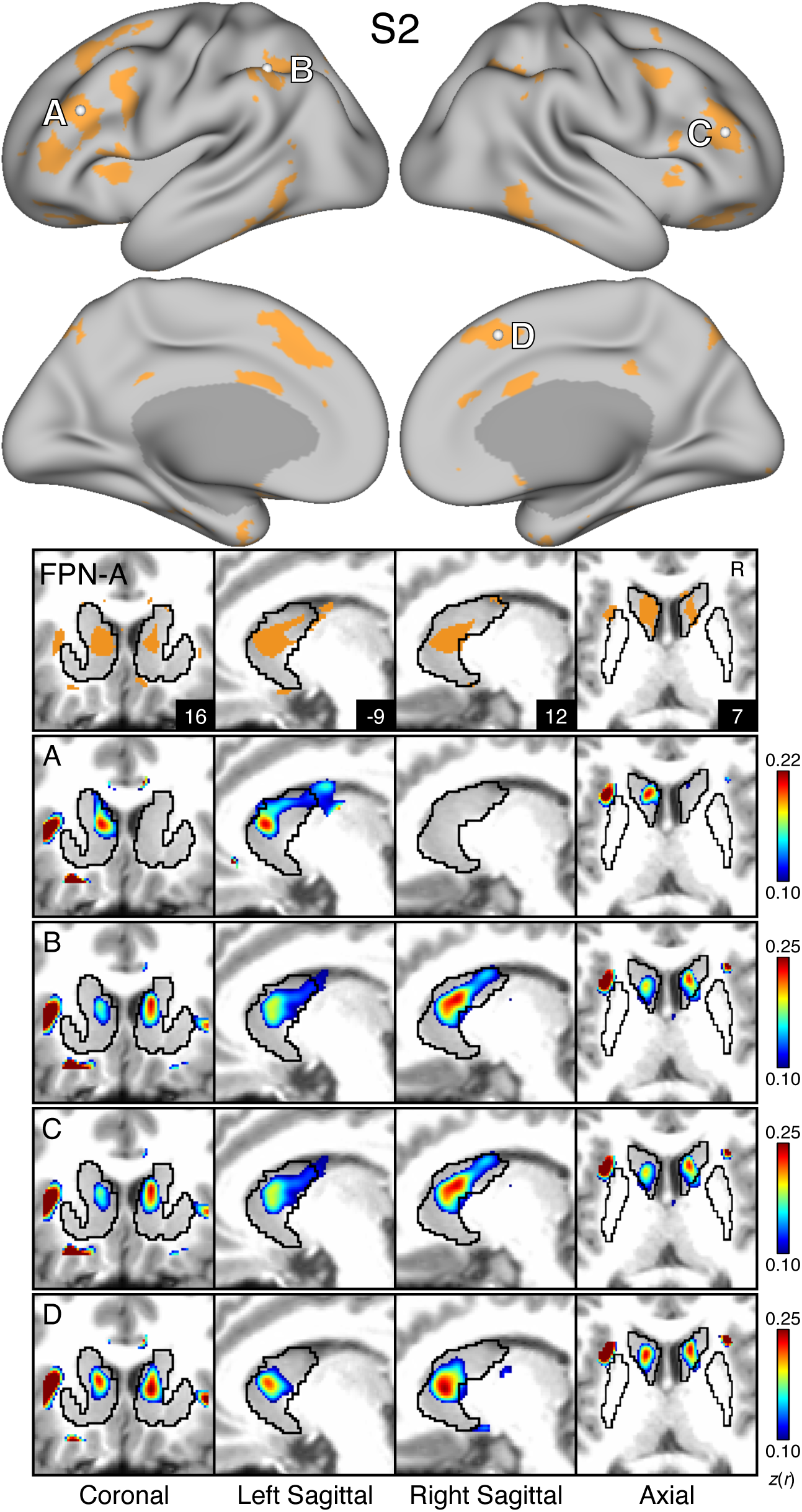
Model-free seed regions in cortical association networks recapitulate Striatal Association Megaclusters. For S1 and S2, the MS-HBM parcellation for a given network is shown on the cortex and in the striatum with the striatal boundaries outlined in black. Four seeds were placed in anterior and posterior nodes (A-D) of cortical association networks FPN-A (orange; Figures S5 and S10), FPN-B (yellow; Figures S6 and S11), LANG (blue; Figures S7 and S12), DN-B (red; Figures S8 and S13), and DN-A (dark red; Figures S9 and S14). Black outlines indicate the boundaries of corresponding individual-specific parcellation-defined networks estimated from the MS-HBM. The correlation maps are plotted as *z*(*r*) with the colorscale at the bottom. There is strong agreement between the seed-based correlation maps and the estimated network parcellation in the striatum.

**Figure S11.**
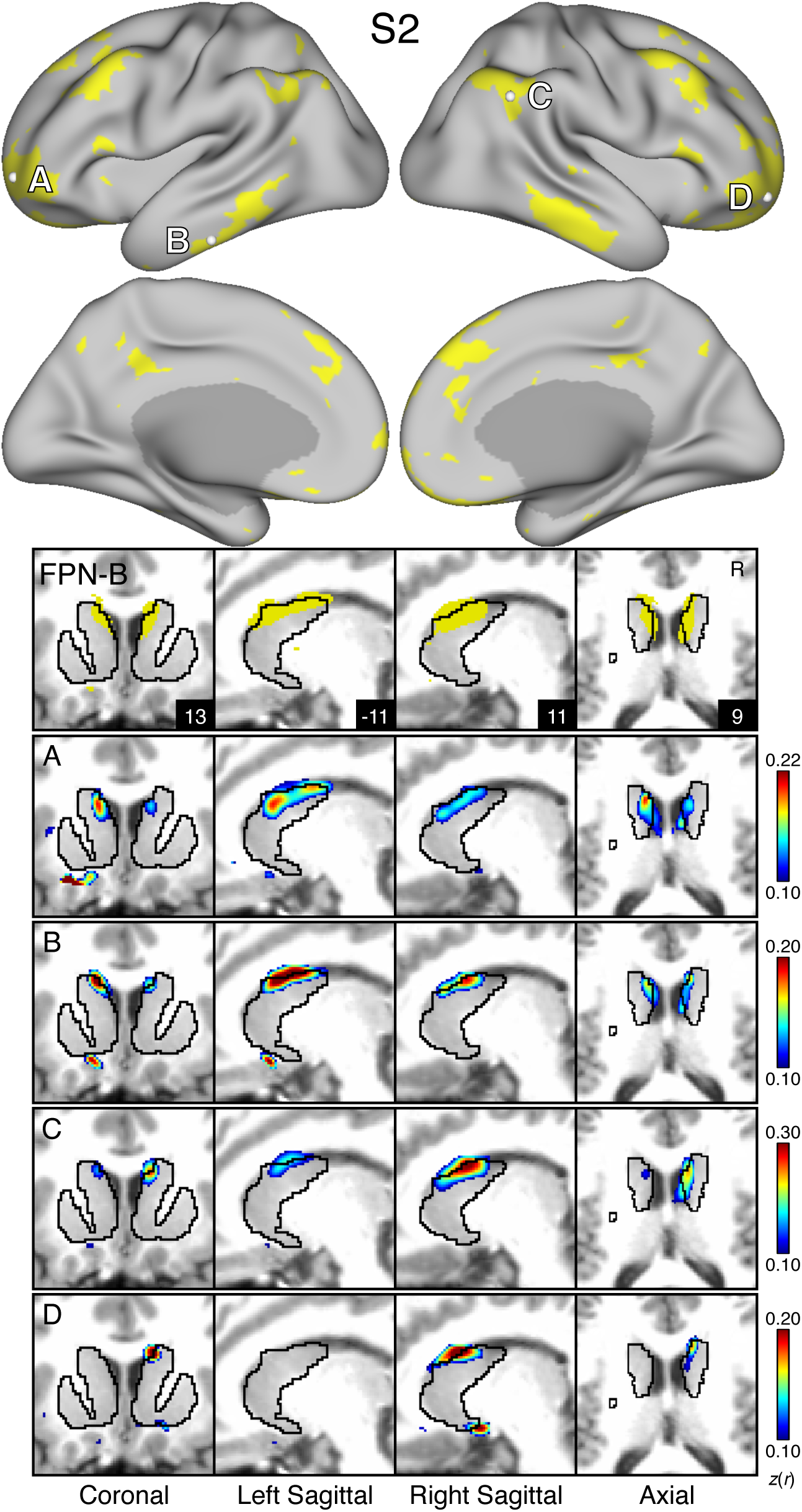
Model-free seed regions in cortical association networks recapitulate Striatal Association Megaclusters. For S1 and S2, the MS-HBM parcellation for a given network is shown on the cortex and in the striatum with the striatal boundaries outlined in black. Four seeds were placed in anterior and posterior nodes (A-D) of cortical association networks FPN-A (orange; Figures S5 and S10), FPN-B (yellow; Figures S6 and S11), LANG (blue; Figures S7 and S12), DN-B (red; Figures S8 and S13), and DN-A (dark red; Figures S9 and S14). Black outlines indicate the boundaries of corresponding individual-specific parcellation-defined networks estimated from the MS-HBM. The correlation maps are plotted as *z*(*r*) with the colorscale at the bottom. There is strong agreement between the seed-based correlation maps and the estimated network parcellation in the striatum.

**Figure S12.**
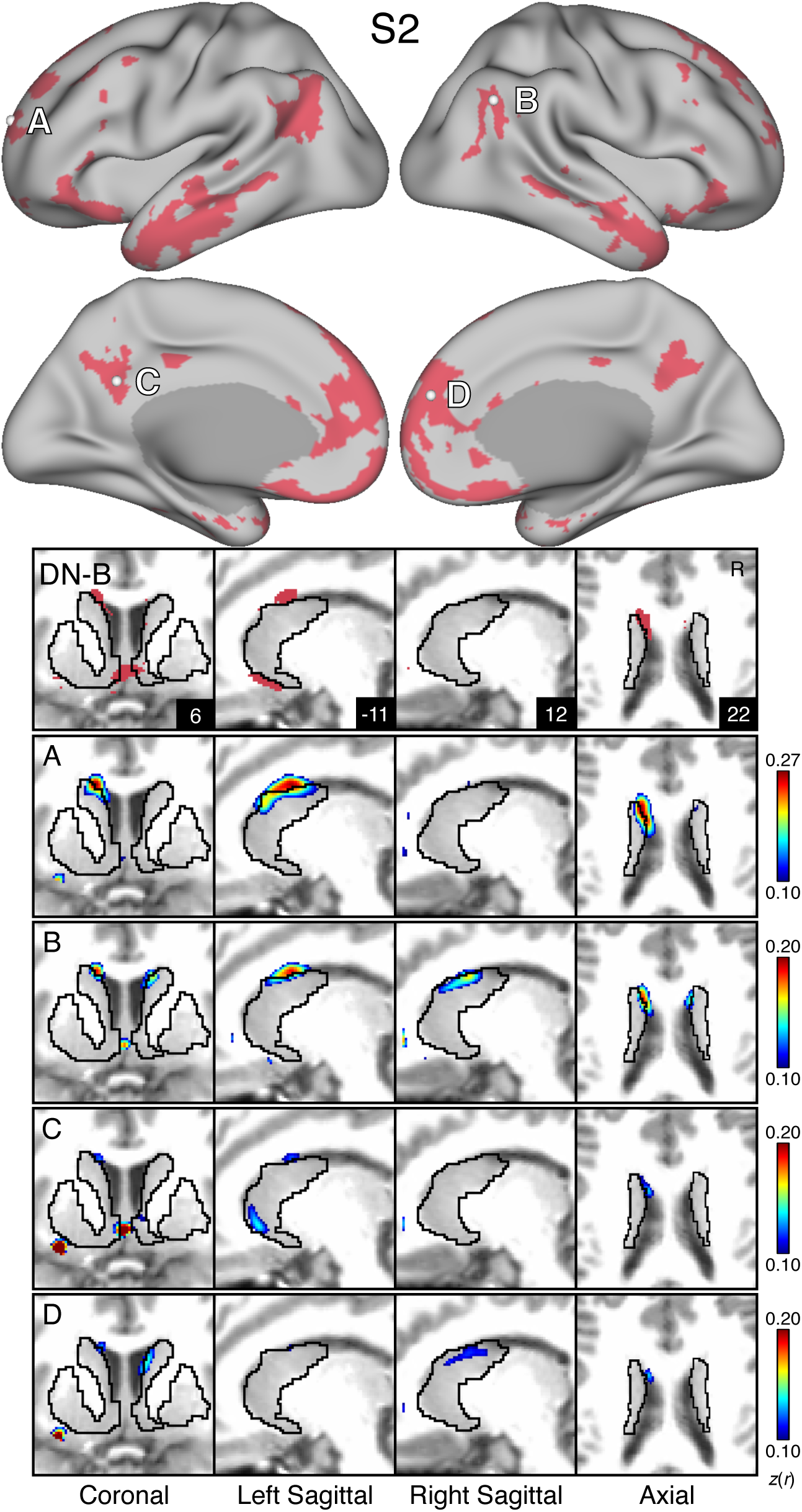
Model-free seed regions in cortical association networks recapitulate Striatal Association Megaclusters. For S1 and S2, the MS-HBM parcellation for a given network is shown on the cortex and in the striatum with the striatal boundaries outlined in black. Four seeds were placed in anterior and posterior nodes (A-D) of cortical association networks FPN-A (orange; Figures S5 and S10), FPN-B (yellow; Figures S6 and S11), LANG (blue; Figures S7 and S12), DN-B (red; Figures S8 and S13), and DN-A (dark red; Figures S9 and S14). Black outlines indicate the boundaries of corresponding individual-specific parcellation-defined networks estimated from the MS-HBM. The correlation maps are plotted as *z*(*r*) with the colorscale at the bottom. There is strong agreement between the seed-based correlation maps and the estimated network parcellation in the striatum.

**Figure S13.**
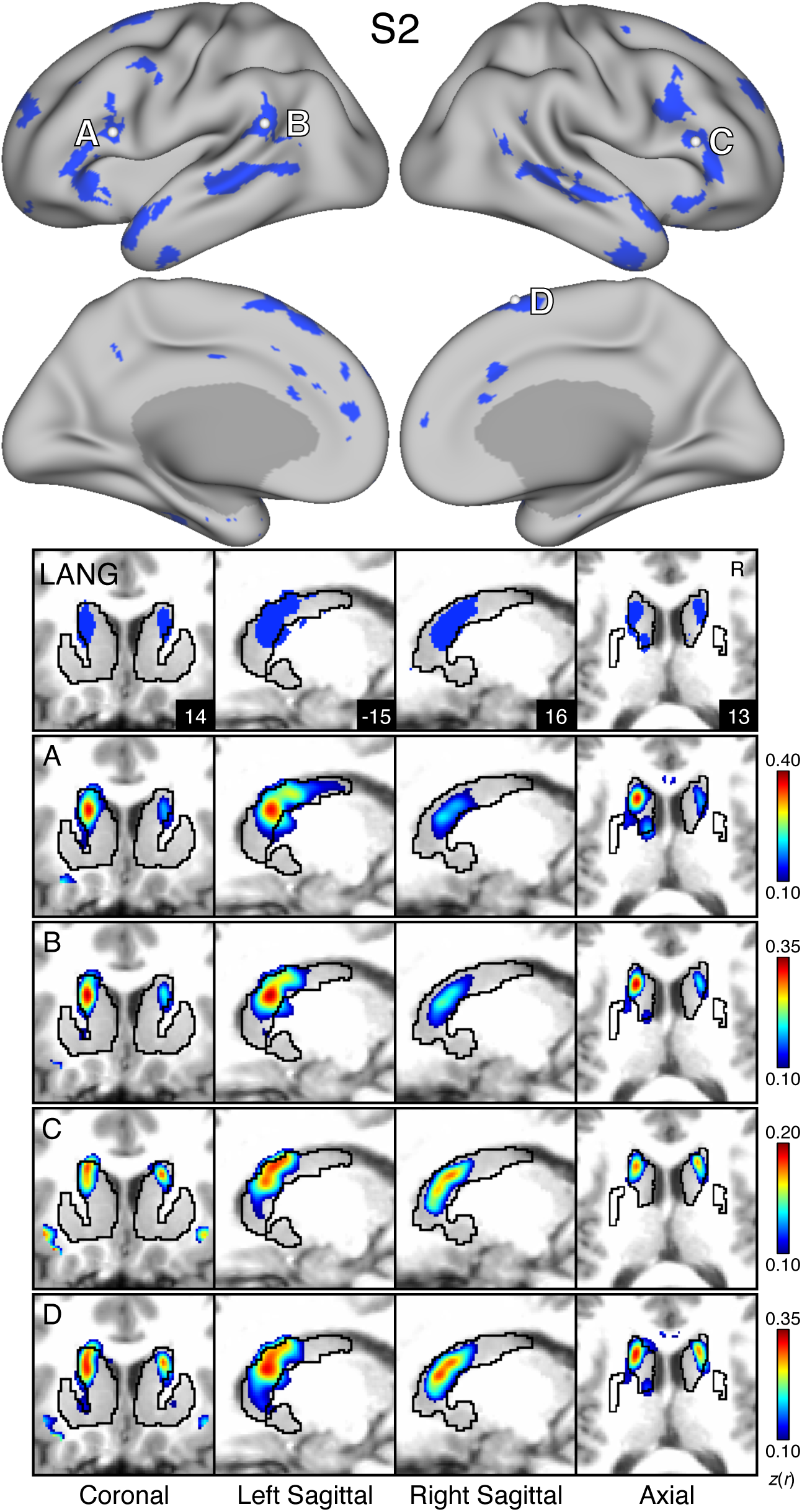
Model-free seed regions in cortical association networks recapitulate Striatal Association Megaclusters. For S1 and S2, the MS-HBM parcellation for a given network is shown on the cortex and in the striatum with the striatal boundaries outlined in black. Four seeds were placed in anterior and posterior nodes (A-D) of cortical association networks FPN-A (orange; Figures S5 and S10), FPN-B (yellow; Figures S6 and S11), LANG (blue; Figures S7 and S12), DN-B (red; Figures S8 and S13), and DN-A (dark red; Figures S9 and S14). Black outlines indicate the boundaries of corresponding individual-specific parcellation-defined networks estimated from the MS-HBM. The correlation maps are plotted as *z*(*r*) with the colorscale at the bottom. There is strong agreement between the seed-based correlation maps and the estimated network parcellation in the striatum.

**Figure S14.**
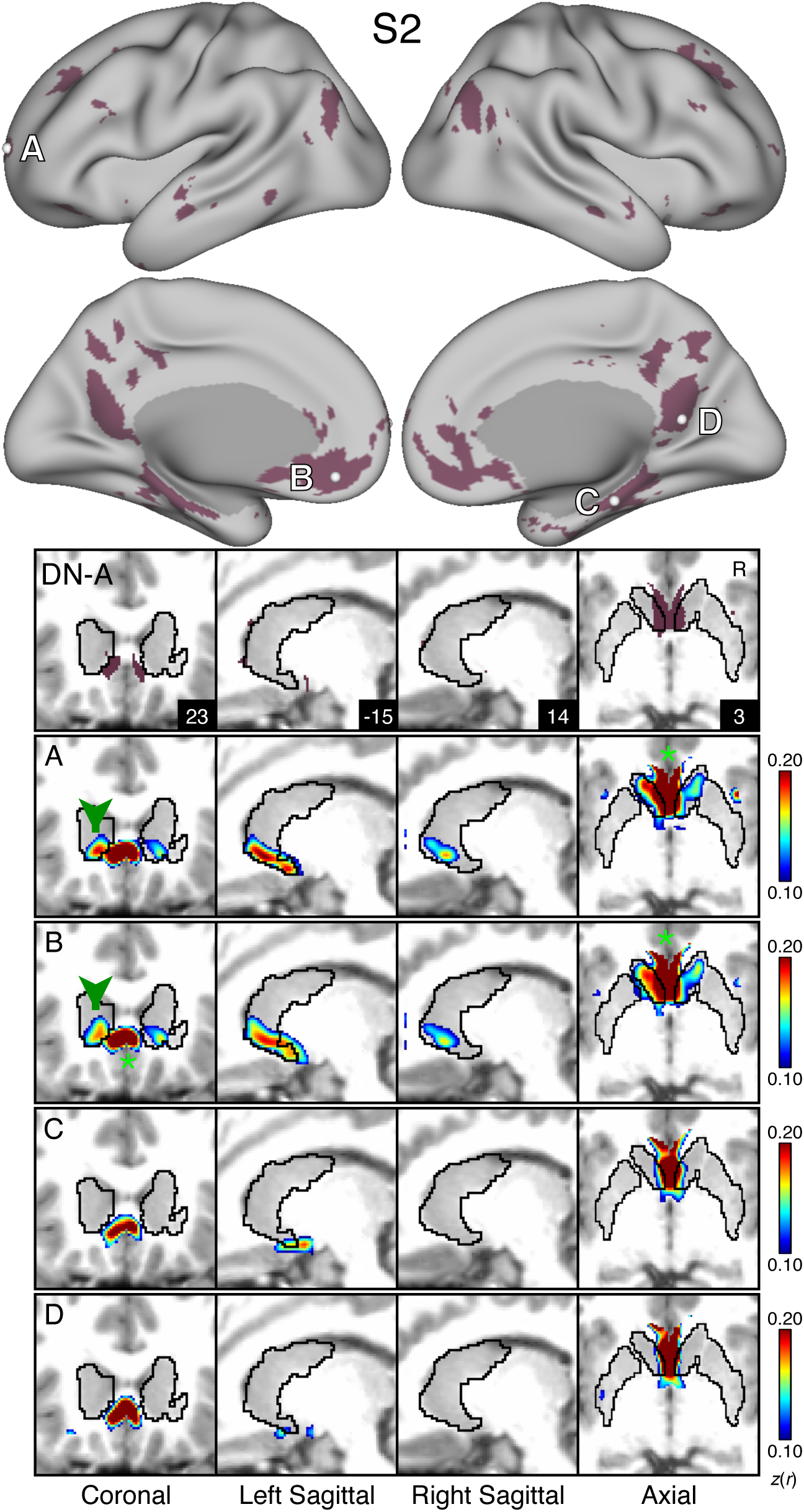
Model-free seed regions in cortical association networks recapitulate Striatal Association Megaclusters. For S1 and S2, the MS-HBM parcellation for a given network is shown on the cortex and in the striatum with the striatal boundaries outlined in black. Four seeds were placed in anterior and posterior nodes (A-D) of cortical association networks FPN-A (orange; Figures S5 and S10), FPN-B (yellow; Figures S6 and S11), LANG (blue; Figures S7 and S12), DN-B (red; Figures S8 and S13), and DN-A (dark red; Figures S9 and S14). Black outlines indicate the boundaries of corresponding individual-specific parcellation-defined networks estimated from the MS-HBM. The correlation maps are plotted as *z*(*r*) with the colorscale at the bottom. There is strong agreement between the seed-based correlation maps and the estimated network parcellation in the striatum.

**Figure S15.**
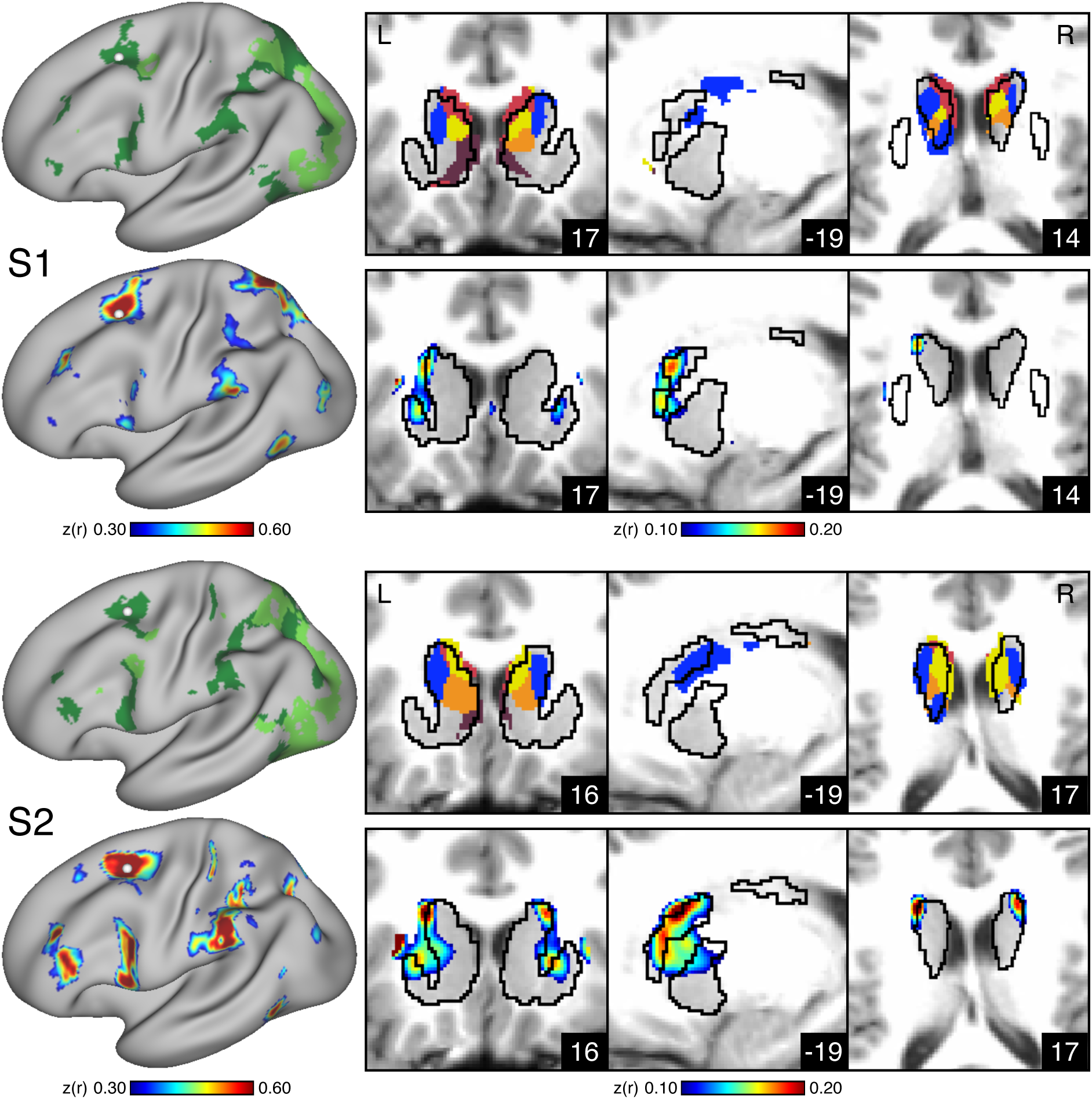
Seed regions in dATN-A correlate with a striatal region lateral to Striatal Association Megaclusters. The frontal eye fields (FEF) and supplementary eye fields (SEF) are cortical regions in dATN-A that project to the caudate in macaques. To test if dATN-A has correlations in the striatum that overlap with Striatum Association Megaclusters, we placed a seed in the approximate location of FEF. Specifically, we used the dATN-A region situated anterior to the precentral gyrus to guide seed placement for each subject. Then, we visualized correlations relative to Striatal Association Megaclusters. The top row for each subject shows the parcellation of dATN-A (green) and dATN-B (dark green) on the left. The seed region is visualized with a white circle. The right-hand side of the top row shows the Striatal Association Megacluster parcellation. The second row shows the correlations with the seed region (∼FEF) on the surface (left) and in the striatum (right). The correlation maps are plotted as *z*(*r*) with the colorscale at the bottom.

## Footnotes

1 A challenge in using masks from T1w structural data is that the BOLD data are lower resolu;on and are smoothed, causing the func;onal response to be spa;ally more extensive. The dilated mask allows the full extent of the correla;on paCerns and parcella;on to be observed.

